# Lipid-associated PML domains regulate CCTα, Lipin1 and lipid homeostasis

**DOI:** 10.1101/2020.03.10.986380

**Authors:** Jonghwa Lee, Jayme Salsman, Jason Foster, Graham Dellaire, Neale D. Ridgway

## Abstract

Nuclear LDs (nLDs) originate at the inner nuclear membrane by a mechanism that involves the promyelocytic leukemia (PML) protein. Here we demonstrate that nLDs in oleate-treated U2OS cells are associated with <u>L</u>ipid-<u>A</u>ssociated <u>P</u>ML (LAP) domains that differ from canonical PML nuclear bodies by the relative absence of SUMO1, SP100 and DAXX. nLDs were also enriched in CTP:phosphocholine cytidylyltransferase α (CCTα), the phosphatidic acid phosphatase Lipin1 and diacylglycerol (DAG). High resolution imaging revealed that LAP domains and CCTα occupy distinct polarized regions on nLDs, and that loss of LAP domains in PML knockout U2OS cells reduced the recruitment of CCTα onto nLDs by its amphipathic α-helical M-domain. The association of Lipin1 and DAG with nLDs was also LAP domain-dependent. The disruption of CCTα and Lipin1 localization on nLDs in PML knockout cells resulted in the inhibition of phosphatidylcholine and triacylglycerol synthesis indicating that LAP domains are a unique PML subdomain involved in nLD assembly and regulation of lipid metabolism.

## INTRODUCTION

Lipid droplets (LDs) are cellular organelles composed of a core of triacylglyceride (TAG) and cholesterol ester (CE) surrounded by a monolayer of phospholipids and associated proteins. In response to nutrient and hormonal signals, fatty acids and cholesterol are stored in or released from LDs to provide energy and biosynthetic precursors as well as buffer the cell from fatty acid toxicity (Sztalryd and Brasaemle, 2017). The largest neutral lipid storage depot is in adipocytes but hepatocytes, enterocytes and macrophages also have the capacity for short-term storage and release of fatty acids from LD (Walther and Farese, 2012). Accumulation of LDs in tissues, caused by an imbalance in lipid storage and hydrolysis, is linked to pathological conditions such as hepatosteatosis, obesity and lipodystrophy.

LDs are proposed to form in the endoplasmic reticulum (ER) by a process that requires the coordinated synthesis of TAG and phospholipids (Henne et al., 2018). TAG initially forms a ‘lens’ in the ER bilayer, expands and pinches off into the cytoplasm, and is coated with ER-derived phospholipids, which regulate surface tension and the storage capacity of LDs (Krahmer et al., 2011). Phosphatidylcholine (PC) is the most significant component of the surface monolayer of LDs (Bartz et al., 2007; Tauchi-Sato et al., 2002), and *de novo* PC synthesis by the CDP-choline pathway in the ER is required for LD biogenesis (Aitchison et al., 2015; Krahmer et al., 2011). In mammalian cells, the CDP-choline pathway is regulated by CTP:phosphocholine cytidylyltransferase (CCT) α and β isoforms that are activated by translocation to nuclear and cytoplasmic membranes, respectively, in response to the content of PC and specific lipid activators sch as fatty acids or type II conical-shaped lipids (e.g. diacylglycerol (DAG) or phosphatidylethanolamine (PE)) (Arnold and Cornell, 1996; Arnold et al., 1997; Xie et al., 2004). During LD biogenesis in fatty acid-treated cells (Gehrig et al., 2008; Lagace and Ridgway, 2005) and during adipocyte differentiation (Aitchison et al., 2015), PC synthesis was increased by CCTα translocation from the nucleoplasm to the inner nuclear membrane (INM). In oleate-treated insect cells, ectopically expressed CCTα and the insect homologue CCT1 translocated from the nucleus to the surface of cytoplasmic LDs (cLDs) resulting in increased PC synthesis to facilitate LD expansion and TAG storage (Guo et al., 2008; Krahmer et al., 2011; Payne et al., 2014). Mammalian CCTα can exit the nucleus under some conditions (Gehrig et al., 2009; Northwood et al., 1999) but does not localize to the surface of cLDs in adipocytes and other cultured cells (Aitchison et al., 2015; Haider et al., 2018).

Interestingly, nuclear lipid droplets (nLDs) were recently identified as site for CCTα translocation and activation of PC synthesis in oleate-treated Huh7 hepatoma cells (Ohsaki et al., 2016). nLDs account for ≈10% of the total cellular LD pool in liver sections and hepatocytes, and have a unique lipid and protein composition (Lagrutta et al., 2017; Uzbekov and Roingeard, 2013). In hepatocytes, nLDs can arise from TAG-rich droplets in the ER lumen that are precursors for VLDL assembly and secretion (Soltysik et al., 2019). These luminal LDs migrate into invaginations of the INM termed the type I nucleoplasmic reticulum (NR) and are released as nascent nLDs into the nucleoplasm where they increase in size by fusion with other nLDs or by *de novo* TAG synthesis (Ohsaki et al., 2016; Soltysik et al., 2019). During or after release into the nucleoplasm, nLDs associate with promyelocytic leukemia (PML) protein, in what were at the time referred to as PML nuclear bodies (PML NB) (Ohsaki et al., 2016). PML NB are dynamic subnuclear domains that regulate gene expression and stress responses and are strongly associated with proteins modified by the small ubiquitin modifier 1 (SUMO1) (Ching et al., 2005; Dellaire and Bazett-Jones, 2004; Van Damme et al., 2010), the most prominent being SP100 nuclear antigen and the death associated domain protein DAXX (Ishov et al., 1999; Zuber et al., 1995) However, of the seven known isoforms of PML, only the PML II isoform was associated with nLDs and required for their formation (Ohsaki et al., 2016). Moreover, the presence of CCTα and perilipin3 on nLDs suggests that these lipid-associated PML-containing domains can regulate lipid synthesis (Soltysik et al., 2019), possibly to generate a metabolic signal to accommodate the uptake and storage of excess fatty acids.

Here we sought to examine the role of PML in nLD biogenesis and function by employing CRISPR/Cas9 to knockout the *PML* gene in human U2OS osteosarcoma cells (PML KO). We found that nLDs in oleate-treated U2OS cells have <u>L</u>ipid-Associated <u>P</u>ML (LAP) domains that are deficient in the canonical PML NB proteins SUMO1, SP100 and death-associated protein 6 (DAXX), and enriched in the lipid biosynthetic enzymes CCTα and Lipin1. Results using PML KO cells indicate that LAP domains are a novel nuclear subdomain complex required for nLD assembly and the recruitment of key enzymes for PC and TAG synthesis.

## RESULTS

### nLDs harbour non-canonical lipid-associated PML (LAP) domains

Similar to hepatoma cells, U2OS osteosarcoma cells contain abundant nLDs that are associated with mCherry-PML-II (Ohsaki et al., 2016). As additional evidence of their nucleoplasmic localization, nLDs in oleate-treated U2OS cells have endogenous PML and CCTα on their surface (Supplemental Fig. S1), and PML-positive nLDs were not encapsulated by the nuclear envelope (NE) based on immunostaining for emerin (Supplemental Fig. S2). Thus, U2OS cells offer an alternate and complementary model to investigate the structure and function of nLDs. Initially we investigated how PML structures on nLDs differ from canonical PML NB by immunostaining control and oleate-treated U2OS cells for the PML NB-associated proteins SUMO1, SP100 and DAXX (Fig. 1 and Supplemental Fig. S3 and S4). U2OS cells contain numerous PML NB that are positive for SUMO1, consistent with the essential role of SUMOylation in PML NB assembly and function (Zhong et al., 2000) (Fig. 1A). Oleate treatment caused the loss of SUMO1-positive PML NB puncta and the appearance of PML-positive nLDs with different levels SUMO1 expression (Fig. 1B). When quantified, a weak or non-existent SUMO1 signal was detected in 75% of PML-positive nLDs (Figure 1C). In addition, DAXX and SP100, proteins whose interaction with canonical PML NB is SUMO-dependent (Ishov et al., 1999) or constitutive (Sternsdorf et al., 1997), respectively, were strongly localized to PML NB in untreated cells but absent or reduced in 80% of PML-positive nLDs in oleate-treated cells (Fig. 1C, Supplemental Fig. S3 and S4). Since the PML structures on nLDs are part of a large lipid complex and relatively devoid of canonical PML NB proteins, we propose that they be designated as <u>L</u>ipid-<u>A</u>ssociated <u>P</u>ML (LAP) domains.

**Figure 1.**
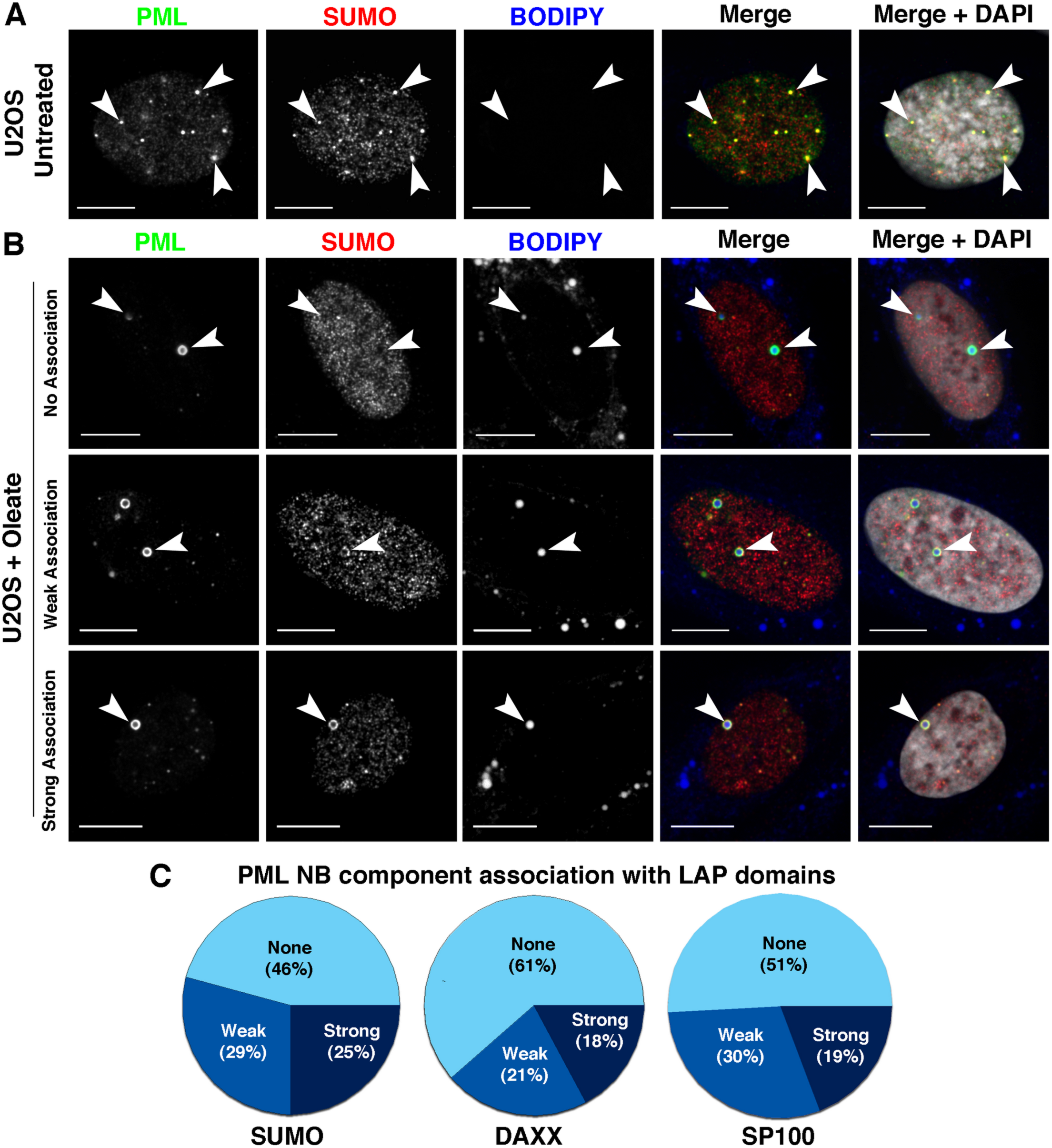
LAP domains on nLDs are deficient in PML NB resident proteins. A and B, U2OS cells were untreated (panel A) or incubated with oleate (400 μM) for 24 h (panel B) prior to immunostaining for PML (green) and SUMO1 (red) and imaging by confocal microscopy. Examples of SUMO1 association (none, weak or strong) with PML-positive nLDs are shown as indicated (panel B). LDs and nuclei were visualized with BODIPY 493/503 and DAPI, respectively (bar, 10μm). C, The intensity of SUMO1, DAXX and SP100 association with PML-positive nLDs was scored as none, weak or strong and determined for at least 100-150 nLDs from at least 50-100 cells. Examples of DAXX and SP100 localization with PML-positive nLDs are shown in Supplemental Fig. S3 and S4.

To investigate how LAP domains influence the lipid and protein composition and biogenesis of nLDs, we utilized U2OS cells in which the *PML* gene was knocked out (PML KO) by CRISPR/Cas9 gene editing (Fig. 2A) (Attwood et al., 2019). In oleate-treated U2OS cells, CCTα was expressed in the nucleoplasm and on the surface of nLDs (Fig. 2B); whereas, in PML KO cells there were fewer BODIPY- and CCTα-positive nLDs, and CCTα was partially localize to the NE. Compared with control cells, the number of cLDs in PML KO cells was reduced slightly (Fig. 2C). However, PML KO cells had a significant reduction in the number of nLD per cell (Fig. 2D) as well as CCTα-positive nLDs (Fig. 2E). The cross-sectional area of cLDs in PML KO cells was similar to controls (Fig. 1F) but there was a significant shift in the distribution toward small nLDs in PML KO cells (Fig. 2G) that was reflected in a 40% decrease in average area (Fig. 2H). CCTα-positive nLDs in control and PML KO cells were similar in size suggesting the enzyme preferentially associates with larger nLDs that are formed by a PML-independent mechanism.

**Figure 2.**
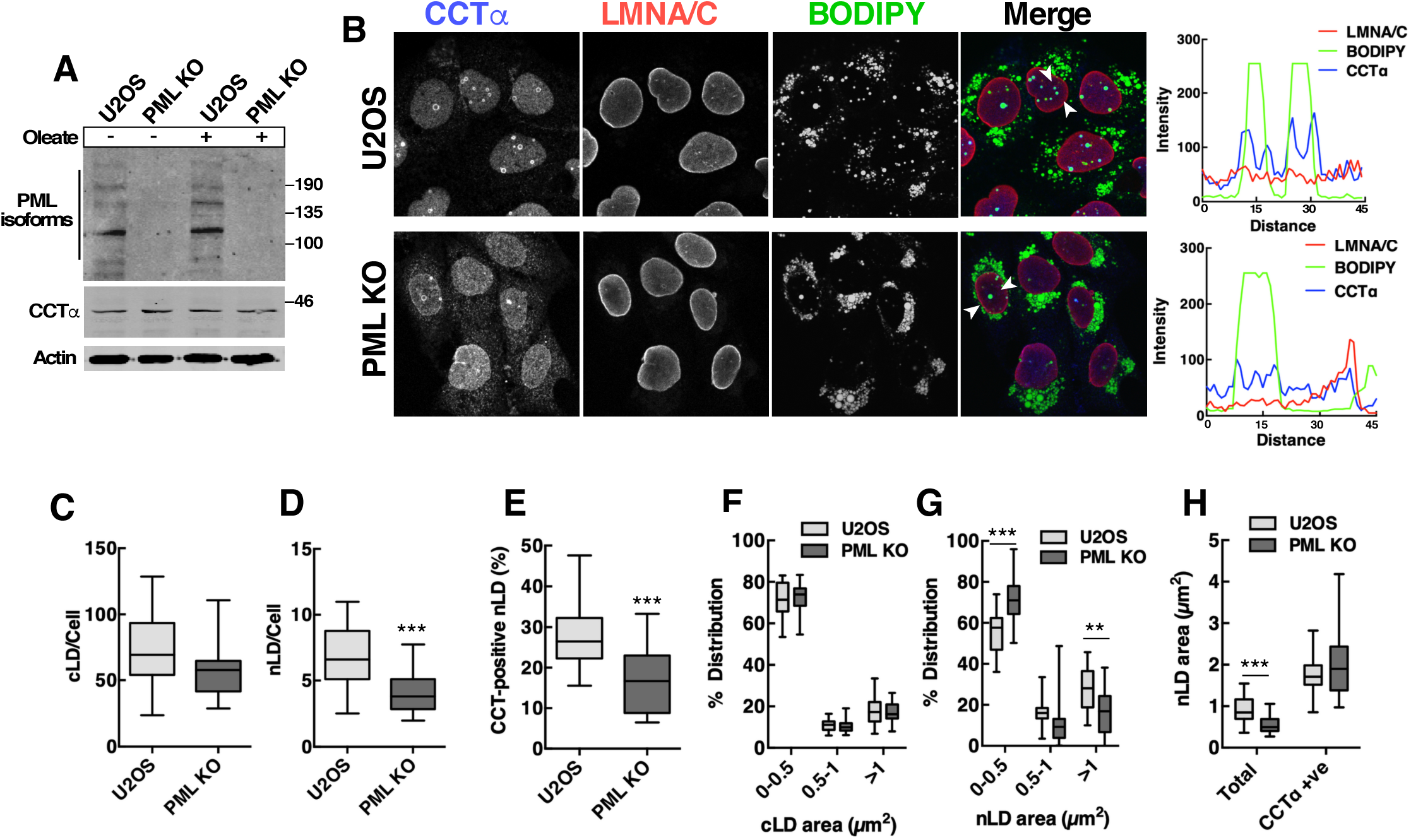
LAP domains regulate biogenesis of nLDs and association of CCTα. A, Lysates of U2OS and PML KO cell that were untreated or incubated with oleate (400 μM) for 24 h were immunoblotted with PML and CCTα-specific antibodies. B, U2OS and PML KO cells were immunostained for CCTα and LMNA/C, and LDs were visualized with BODIPY 493/503. Arrows indicate the region for RGB line graphs showing the localization of CCTα on the surface of nLDs. C and D, quantitation of cytoplasmic and nuclear LDs in cells treated with oleate (400 μM) for 24 h. E, quantitation of CCTα-positive nLDs in oleate-treated cells. F and G, the size distribution of cLDs (panel F) and nLDs (panel G) in oleate-treated cells was determined by cross-sectional area binning. H, average cross-sectional area of total and CCTα-positive nLDs was quantified in oleate-treated cells. In panels C-H, results are presented as box and whisker plots showing the mean and 5^th^-to-95^th^ percentile for analysis of 50-100 cells from 3 separate experiments. Significance was determined by two-tailed t-test compared to matched U2OS cell controls (\*\**p*<0.01; \*\*\**p*<0.001).

To identify the PML isoform involved in nLD formation in U2OS cells, PML KO cells were individually transfected with GFP-tagged version of the 7 PML isoforms. In agreement with results in Huh7 cells (Ohsaki et al., 2016), GFP-PML-II associated with nLDs and increased their abundance in PML KO cells while other PML isoforms did not associate or correct the PML KO phenotype (Supplemental Fig. S5A and B, results not shown). GFP-PML-II expression in PML KO cells significantly increased the number and size of nLDs to the level observed in U2OS cells (Supplemental Fig. S5C and E) and also restored the slight reduction in parameters for cLDs (Supplemental Fig. S5D and F). The specific requirement for PML-II in nLD formation is further evidence that LAPs are a unique nuclear PML subdomain.

### CCTα and LAP domains localize to discrete regions of nLDs

CCTα associates with membranes via its amphipathic α-helical M-domain, which is antagonized by phosphorylation of 16 serine, threonine and tyrosine residues in the adjacent P-domain (Fig. 3A) (Cornell and Ridgway, 2015). The binding of CCTα to LD involves large hydrophobic side-chains in M-domain that insert into voids in the phospholipid monolayer (Prevost et al., 2018). To determine if the M- and P-domains regulate binding to nLDs, CCTα with truncations and point mutations in these domains was expressed in U2OS cells (Fig. 3B) and localization on nLDs was quantified by immunofluorescence microscopy (Fig. 3C and D). CCTα localization to nLDs was completely prevented by mutation of 8 lysine residues in the M domain (CCTα-8KQ) that form electrostatic interactions with membrane lipids (Johnson et al., 2003). Conversely, CCTα-3EQ, a M-domain mutant with enhanced membrane association (Johnson et al., 2003), was localized to nLDs as well as the NE. Deletion of the P-domain (CCTα-ΔP) and a dephosphorylated mimic with 16 serine residues mutated to alanine (CCTα-16SA) were strongly associated with nLDs, whereas the phosphorylated mimic with 16 serine residues mutated to glutamate (CCTα-16SE) was not detected on nLDs. Catalytic dead CCTα-K122A was localized on nLDs similarly to wild-type. These results show that interaction of CCTα with the nLD phospholipid monolayer is mediated by electrostatic interactions with the M-domain and antagonized by phosphorylation of the P-domain.

**Figure 3.**
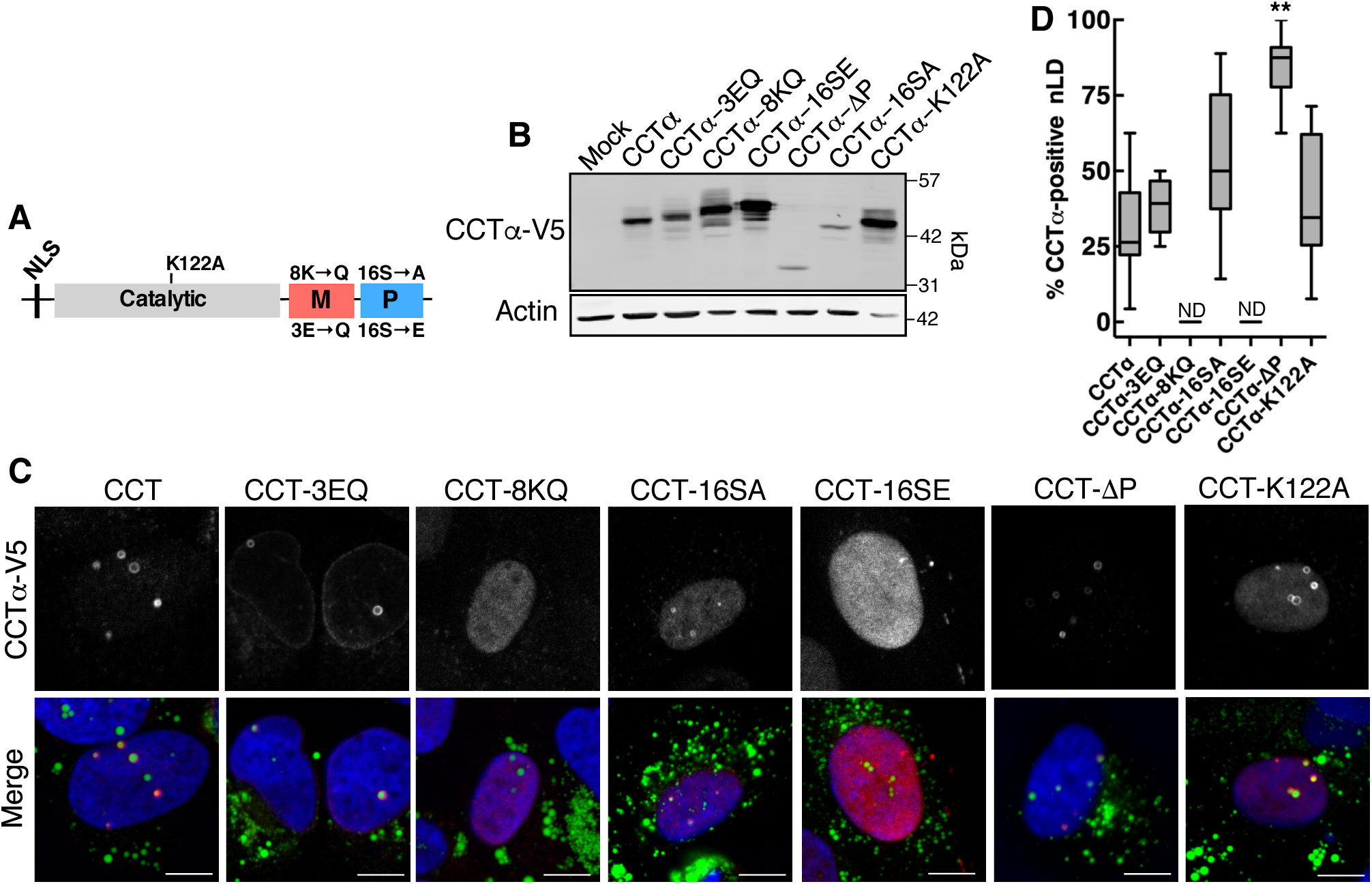
The interaction CCTα with nLDs is dependent on its M- and P-domains. A, Domain organization of CCTα showing the location of catalytic, M- and P-domain mutations. B, U2OS cells were transiently transfected with empty vector (Mock) or the indicated CCTα−V5 constructs and treated with oleate (400 μM) for 24 h. Total cell lysates were immunoblotted with V5 and actin antibodies. C, Confocal images of U2OS cells (transfected and treated as described in panel B) immunostained with a V5 antibody. LDs and nuclei were visualized with BODIPY 493/503 and propidium iodide, respectively (bar 10 μm). D, Quantification of the percentage of nLDs that were positive for wild-type CCTα-V5 mutants presented as box and whisker plots showing the mean and 5^th^-to-95^th^ percentile from analysis of 10-20 fields of cells from 2-3 separate experiments. Significance was determined by one-way ANOVA and Tukey’s multiple comparison to CCTα- V5 (ND, not detected; \*\**p*<0.01).

Next we used spinning disk confocal 3D and super-resolution radial fluctuation (SRRF) (Gustafsson et al., 2016) imaging to determine how endogenous CCTα and LAP domains are disposed on the surface of nLDs (Fig. 4). 3D reconstruction of a typical nLD revealed a lipid body (BODIPY) located close to the basal surface of the INM (Fig. 4A). CCTα coated most of the nLD but was absent from the top of the particle (Fig. 4A, d), which was occupied by a ‘cap’ of PML protein (Fig. 4A, e). SRRF imaging of this nLD in a cross-section where CCTα and PML intersect (indicted by the yellow line in Fig. 4A, b-f) revealed a punctate distribution (Fig. 4B and C) consistent with each protein occupying non-overlapping, interdigitated regions on the nLD surface (error and resolution analysis using NanoJ-SQUIRREL (Culley et al., 2018) for this image is shown in Supplemental Fig. S6). Additional 3D renderings of 8 nLD are shown in Supplemental Fig. S7. In the case of smaller nLDs (Supplemental Fig. S7A1, C5 and C6), PML formed a patch on the surface that minimally overlapped with CCTα. Similar to Fig. 4, another large nLD displayed a polarized PML patch (Supplemental SFig. 7B1). More infrequent were large nLD with PML coating much of the surface and reduced CCTα association (Supplemental SFig. S7B4). In summary, LAP domains generally occupy only part of the surface of an nLD and have limited overlap with CCTα, which more strongly associates with large nLDs via its M-domain.

**Figure 4.**
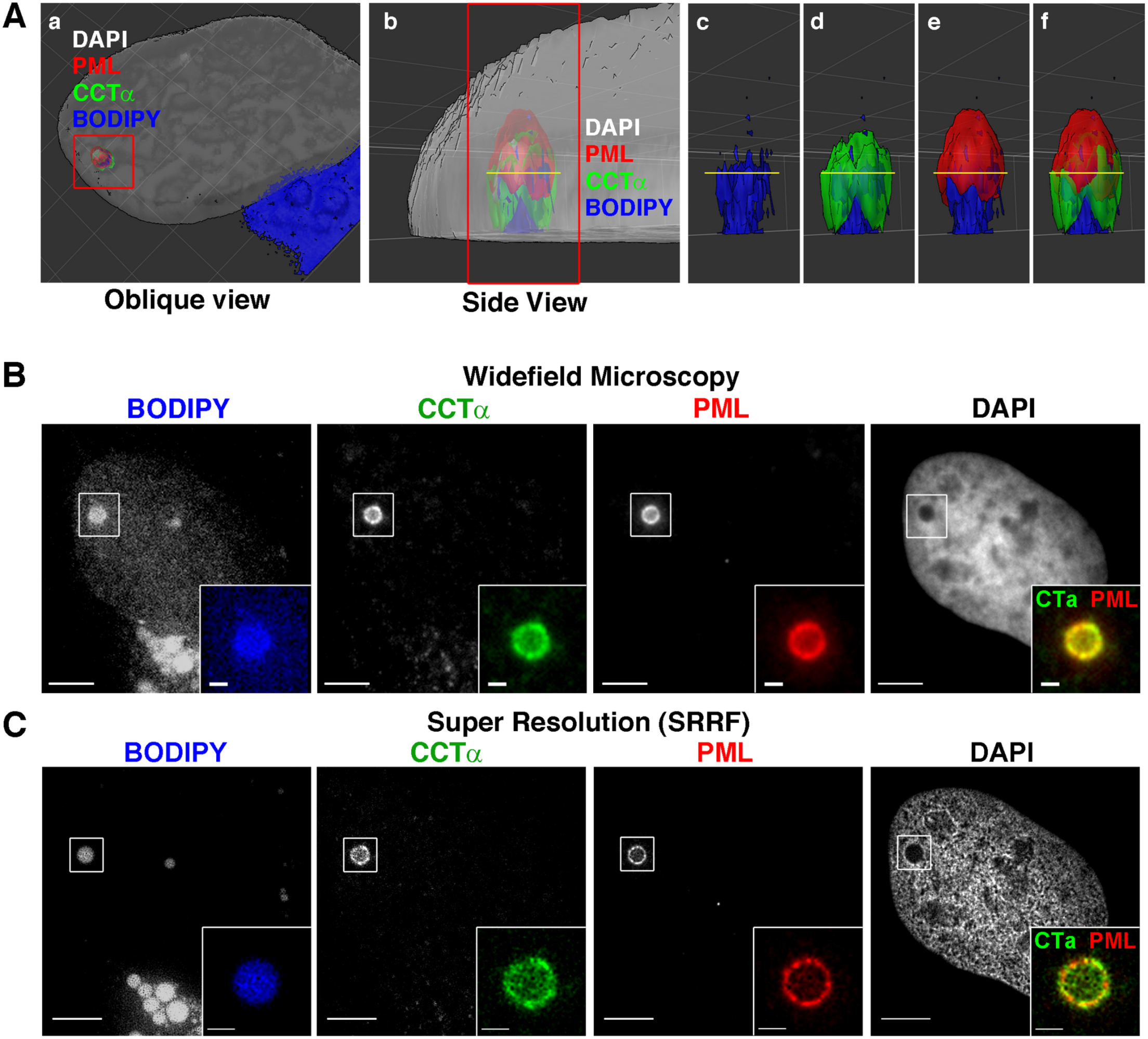
High resolution imaging of LAP domains and CCTα on nLDs. A, 3D surface reconstruction of image stacks of oleate-treated (400 μM, 24 h) U2OS cells immunostained for PML and CCTα. LDs and nuclei were visualized with BODIPY 493/503 and DAPI, respectively. A large nLD was identified in association with CCTα and PML (panel a, red frame). The image was rotated and zoomed to produce a side view of the nLD (panel b, red frame). The framed structure in b was assessed by removing the DAPI channel (panels c-f) to reveal how the nLD/BODIPY (blue), CCTα (green) and PML (red) are associated. B, Widefield images of the same nLD in A imaged at the level of the yellow line indicated in panels b-f (image rotated ∼90° clockwise). The nLD is highlighted and magnified in the inset (bar, 5µm; Inset scale bar, 1µm). C, Images of the same field of view in B were acquired by NanoJ SRRF. The nLD is highlighted and magnified in the inset (bar, 5µm; inset bar, 1µm).

### The DAG and Lipin1 content of nLDs is regulated by LAP domains

In addition to being a precursor for the TAG and PC that is incorporated into LD, DAG also stimulates translocation and activation of CCTα on membranes (Arnold et al., 1997). Thus, we investigated whether CCTα interaction with nLDs could be regulated by DAG as a mechanism to coordinate PC and TAG synthesis. Initially, we visualized DAG in cells cultured with or without oleate for 24 h using the DAG biosensor GFP-C1(2)δ (Codazzi et al., 2001) (Fig. 5). In untreated U2OS and PML KO cells, the DAG biosensor was localized on reticular and punctate structures in the cytoplasm (Fig. 5A). Exposure of U2OS cells to oleate resulted in the appearance of the DAG biosensor on dispersed cytoplasmic structures that did not correspond to cLDs, and occasionally on LipidTox Red-positive nLDs (Fig. 5B, a-c). In contrast, the DAG biosensor was strongly associated with cLDs in PML KO cells and less frequently observed on nLDs (Fig. 5C, a-c). This is the first evidence that PML regulates the DAG content of cLDs and nLDs.

**Figure 5.**
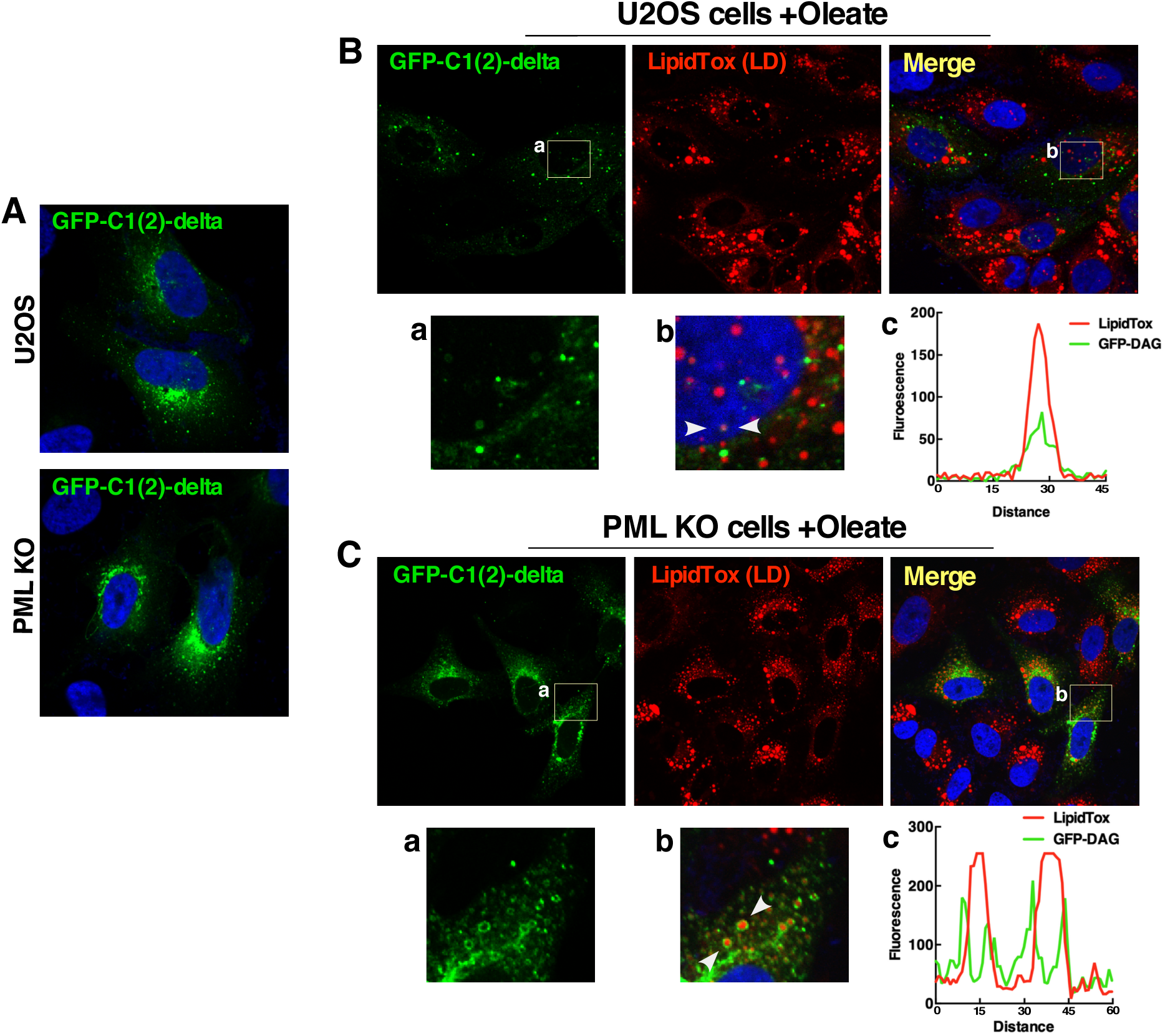
LAP domains control DAG levels on cLDs and nLDs. A, Localization of the DAG sensor GFP-C1(2)δ in U2OS and PKL KO cells. B, U2OS cells expressing GFP-C1(2)δ were treated with oleate (400 μM) for 24 h and LDs were visualized with LipidTox Red. Highlighted areas from GFP and merged images are shown in panels a and b. Arrows indicate the region selected for an RGB line plot (c) showing the association of the DAG sensor with a nLD. B, PML KO cells were treated as described above. Highlighted areas from GFP and merge images are shown in panels a and b. Arrows indicate the region selected for an RGB line plot (c) showing the association of the DAG sensor with cytoplasmic LDs.

To better assess how PML regulated the levels of DAG on nLDs, a tandem nuclear localization signal (NLS) was inserted in GFP-C1(2)δ (nGFP-DAG). When transiently expressed in oleate-treated U2OS cells, the nuclear biosensor nGFP-DAG was detected on two types of structures; small puncta that did not stain with LipidTox Red, and LAP-positive nLDs (Fig. 6A). For the purpose of quantification, we divided LipidTox Red-positive nLDs in U2OS cells into four populations based on the presence or absence of DAG and/or PML (Fig. 6B). This revealed that 60% of total nLDs were negative for both DAG and PML (DAG/PML-). The majority of the remaining nLDs contained DAG and PML (79%), with the remainder containing only DAG (7%) or PML (14%) (Fig. 6B, insert). The cross-sectional area of DAG/PML- nLDs was significantly reduced compared to total nLDs, while the PML+ and DAG/PML+ nLDs were significantly larger (Fig. 6C). DAG+ nLDs were virtually absent from PML KO cells compared to total and DAG-negative nLDs, both of which were also significantly reduced (Fig. 6D). The nuclear sensor also detected DAG on the abundant nLDs that form in oleate-treated Caco2 cells (Supplemental Fig. S8A). These data indicate that DAG is primarily enriched in large LAP-positive nLDs, and that PML KO reduces DAG in nLDs but increases DAG in cLDs.

**Figure 6.**
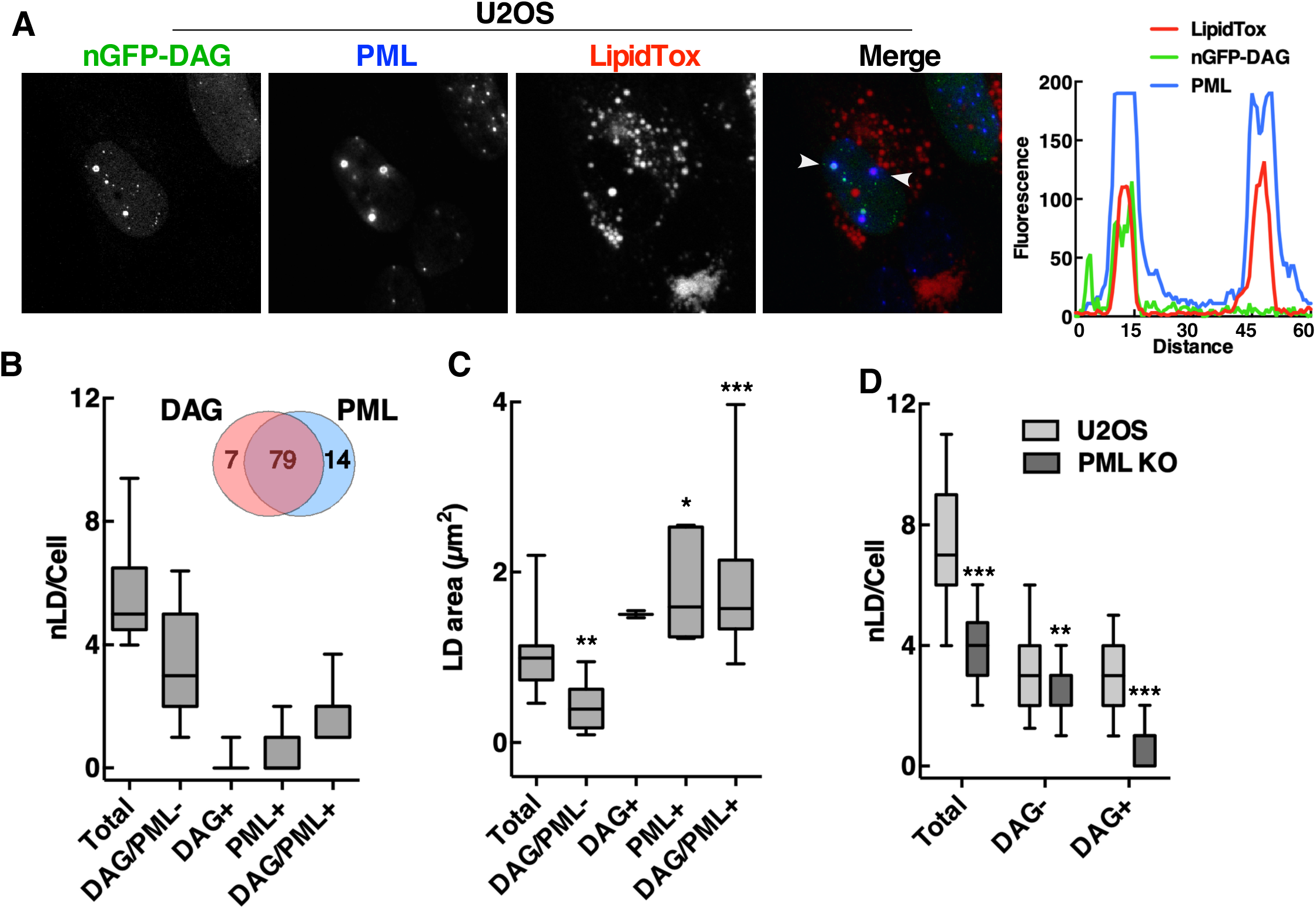
LAP-positive nLDs are enriched in DAG. A, U2OS cells transiently expressing nuclear localized GFP-C1(2)δ (DAG-GFP) were exposed to oleate (400 μM) for 24 h were fixed and immunostained with a PML antibody. LDs were visualized with LipidTox Red. Arrows indicate the region selected of a RGB line plot showing the localization of DAG-GFP with PML and nLDs. B, Quantitation of nLDs in oleate-treated U2OS cells containing neither DAG-GFP or PML (DAG/PML-), DAG-GFP (DAG+), PML (PML+) or both (DAG/PML+). A Venn diagram shows the percent distribution of DAG-GFP and PML positive nLDs. C, the average cross-sectional area of nLDs in oleate-treated U2OS cells containing DAG-GFP and/or PML as described above. D, Quantification of DAG-negative (+) and DAG-negative (-) nLDs in U2OS and PML KO cells. Results in panels B, C and D are presented as box and whisker plots showing the mean and 5^th^-to- 95^th^ percentile from analysis of 50-100 cells in 3 separate experiments. Significance was determined by one-way ANOVA and Tukey’s multiple comparison to total nLD area (C) or two-tailed t-test compared to U2OS controls (D) (\**p*<0.05; \*\**p*<0.01; \*\*\**p*<0.001).

To determine whether DAG was a positive effector of CCTα association with nLDs, we determined whether CCTα was preferentially enriched in DAG-positive nLDs. Immunofluorescence and RGB plots through nLDs in U2OS cells showed association of the nuclear DAG sensor and CCTα (Fig. 7A). In PML KO cells, there was evidence of CCTα-positive nLDs that lacked DAG suggesting poor correlation between DAG content and CCTα, a conclusion supported by quantitation of CCTα and DAG distribution on nLDs. Of the approximately 50% of nLDs in U2OS cells that contained DAG and/or CCTα, only 43% contained both DAG and CCTα suggesting that the DAG content of an nLDs is not a prerequisite for recruitment of CCTα (Fig. 7B, insert). Quantification of DAG/CCTα distribution on nLDs in PML KO cells indicated a reduction in all four populations compared to control cells, particularly those that contained DAG and DAG/CCTα but the percent distribution of DAG and CCTα on nLDs was similar to control cells (Fig. 7B, insert). As well, there was no significant difference in the cross sectional area of nLDs containing DAG/CCTα in control versus PML KO cells (Fig. 7C). Collectively this shows that the presence of CCTα and DAG on nLDs is poorly correlated in both control and PML KO cells, a conclusion that is consistent with charged residues in the CCTα M-domain being a driving force for interaction with nLDs (Fig. 3).

**Figure 7.**
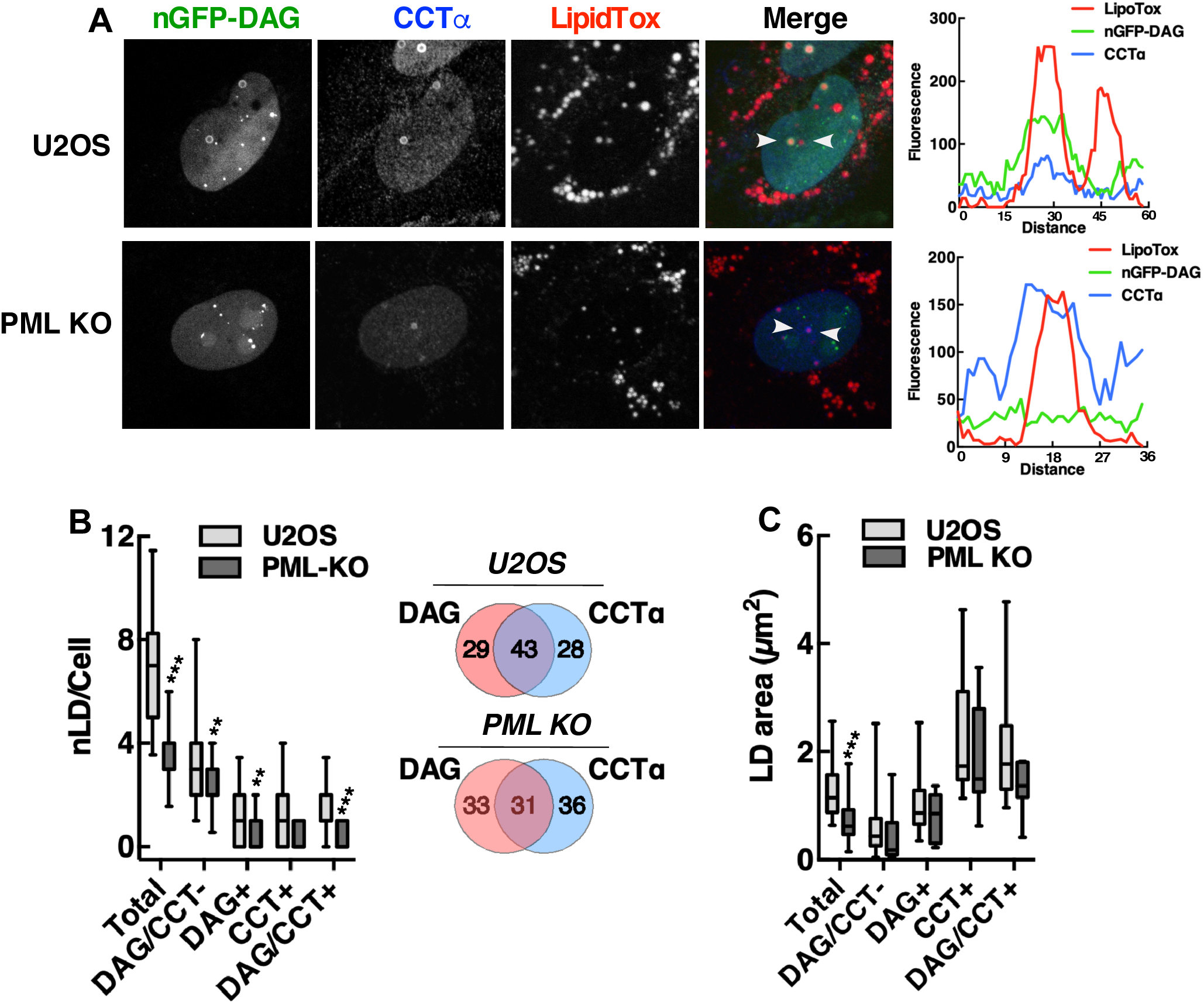
CCTα association with nLDs is DAG independent. A, U2OS and PML KO cells transiently expressing a nuclear DAG-GFP sensor were exposed to oleate (400 μM) for 24h, fixed and immunostained with a CCTα antibody, and incubated with LipidTox Red to visualize LDs. Arrows indicate the region selected for a RGB plots showing the association of CCTα and DAG-GFP. B, Quantification of nLDs in oleate-treated U2OS and PML KO cells containing neither DAG-GFP or CCTα (CCTα/DAG-GFP-), DAG-GFP (DAG+), CCTα (CCT+) or both (CCT/DAG-GFP+). A Venn diagram shows the percent distribution of DAG-GFP and CCTα-positive nLDs. C, in oleate-treated U2OS and PML KO cells, the cross-sectional area of nLDs containing DAG-GFP and/or CCTα was measured. Results in panels B and C are presented as box and whisker plots showing the mean and 5^th^-to-95^th^ percentile for analysis of 50-100 cells from 3 separate experiments. Significance was determined by a two-tailed t-test compared to matched U2OS controls (\*\**p*<0.01; \*\*\**p*<0.001).

The source for DAG in nLDs could be Lipin1, a nuclear/cytoplasmic PA phosphatase that produces DAG for phospholipid and TAG synthesis (Csaki et al., 2013). The human and murine genes encode similar Lipin1α and β splice variants (Croce et al., 2007; Peterfy et al., 2005) that are partially localized to cLDs in COS cells and macrophages (Valdearcos et al., 2011; Wang et al., 2011), potentially recruited by seipin (Sim et al., 2012). To determine if Lipin1 is recruited to nLDs by a LAP-dependent mechanism, transiently expressed Lipin1α and β were localized in U2OS and PML KO cells by immunofluorescence microscopy. V5-tagged Lipin1α was expressed in the nucleus and cytoplasm of untreated U2OS and PML KO cells (Fig. 8A). Treatment of U2OS cells with oleate caused extensive association of Lipin1α with the surface of BODIPY-positive nLDs but not with cLDs (Fig. 8B, a). Lipin1α was also localized to the surface of nLDs in Caco2 cells treated with oleate for 24 h (Supplemental Fig. S8B). To determine the extent of Lipin1α and DAG co-localization in U2OS cells, the distribution and cross-sectional area of Lipin1α- and DAG-containing nLDs was quantified. Of the DAG/Lipin1α-positive nLDs, 53% contained both (Fig. 8C, insert), and Lipin1α-positive nLDs were significantly larger than other components of the population (Fig. 8D). In oleate-treated PML KO cells, Lipin1α was diffusely localized in the cytoplasm and nucleus but was virtually absent from nLDs (Fig. 8E), a conclusion that was confirmed by quantification of Lipin1α-positive nLDs in PML KO cells compared to controls (Fig. 8F). The Lipin1β isoform also associated with the surface of nLDs but not cLD in oleate-treated U2OS cells (Supplemental Fig. S9A), and was not detected on nLDs in oleate-treated PML KO cells (Supplemental Fig. S9B). The Lipin1 substrate PA was detected with the GFP-nes-2xPABP (Bohdanowicz et al., 2013) biosensor on the PM and cytoplasmic membranes of control and PML KO cells but was absent from nLDs or cLDs (Supplemental Fig. S10). These data indicate that LAP domains are required for association of Lipin1 and DAG with nLDs.

**Figure 8.**
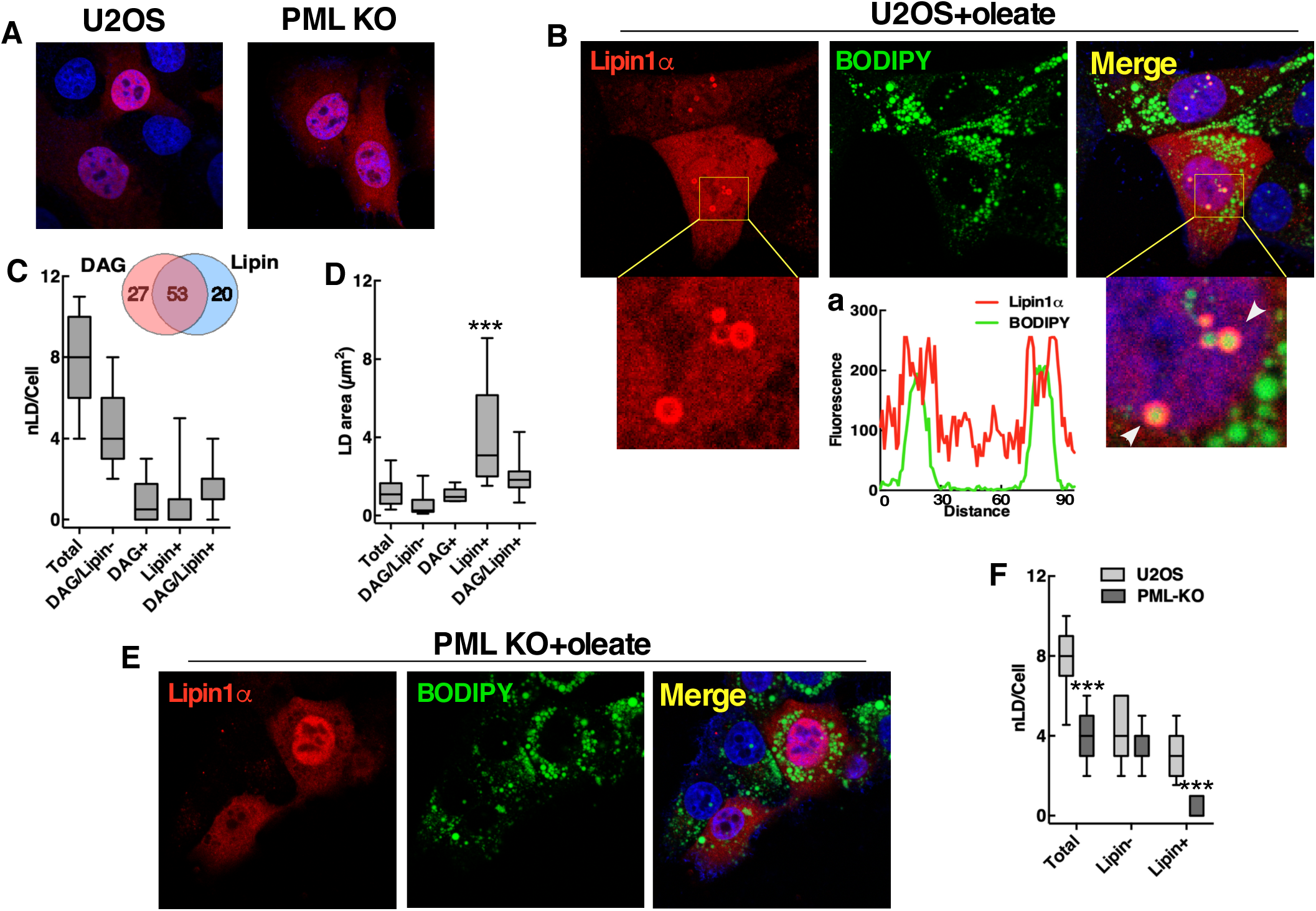
LAP domains promote formation of Lipin1α with DAG-positive nLDs. A, U2OS and PML KO cells transiently expressing Lipin1α-V5 were immunstained with a V5 monoclonal antibody and nuclei were visualized with DAPI. B, U2OS cells transiently expressing Lipin1α-V5 were treated with oleate (400 μM) for 24h and immunostained with a V5 monoclonal antibody. LDs were visualized with BODIPY 493/503. Co-localization of Lipin1α and nLDs is shown in selected regions from Lipin1α (a) and merge (b) images. Arrows show the region selected for a RGB line plots (c) showing Lipin1α on the surface of nLDs. C, Quantification of nLDs in oleate-treated U2OS cells containing neither DAG-GFP or Lipin1α (Lipin/PML-), DAG-GFP (DAG+), Lipin1α (Lipin+) or both (Lipin/DAG+). A Venn diagram shows the percent distribution of DAG-GFP and Lipin1α-positive nLDs. D, The cross-sectional area of nLDs containing differing compositions of DAG-GFP and Lipin1α was quantified in oleate-treated U2OS cells. E, PML KO cells were treated with oleate and immunostained as described above. F, Quantification of Lipin-negative (-) and Lipin-positive (+) nLDs in U2OS and PML KO cells. Results in panels C, D and F are presented as box and whisker plots showing the mean and 5^th^-to-95^th^ percentile for analysis of 50-100 cells from 3 separate experiments. Significance was determined by one-way ANOVA and Tukey’s multiple comparison (panels C and D) or two-tailed t-test comparted to control U2OS cells (panel F) (\*\*\**p*<0.001).

### LAP domains positively regulate PC and TAG synthesis

We next tested whether ablating LAP domains in PML KO cells and reducing nLD-associated CCTα, DAG and Lipin1 affected cellular lipid synthesis. PC biosynthesis was measured by [^3^H]choline pulse-labelling for 2 and 4 h in cells treated with or without oleate. PC synthesis in untreated PML KO cells was similar to controls (Fig. 9A). However, PC synthesis was poorly activated in oleate-treated PML KO cells and reduced significantly compared to the 2.5-fold activation caused by oleate in U2OS cells (Fig. 9A). PML KO had a minor inhibitory effect on *de novo* synthesis of fatty acids from [^3^H]acetate that was only significant in cells cultured in LPDS (Fig. 9B). TAG and CE synthesis in control and PML KO cells was measured by [^3^H]oleate incorporation (Fig. 7E and F). Relative to controls, the incorporation of [^3^H]oleate into TAG and CE in PML KO cells was significantly decreased ∼2-fold. To measure the *de novo* synthesis of TAG, control and PML KO cells were incubated with 100 μM oleate for up to 6 h in the presence of [^3^H]glycerol. In this case, incorporation into TAG in PML KO cells was increased at 1-4 h but returned to control values at 6 h (Fig. 7D). [^3^H]Glycerol incorporation into TAG in U2OS and PML KO cells was similar after oleate treatment for 12 and 24 h (results not shown). Despite accounting for only 10-15% of total cellular LDs, nLDs with associated CCTα and Lipin1 activities are important sites for the regulation of PC and TAG synthesis.

**Figure 9.**
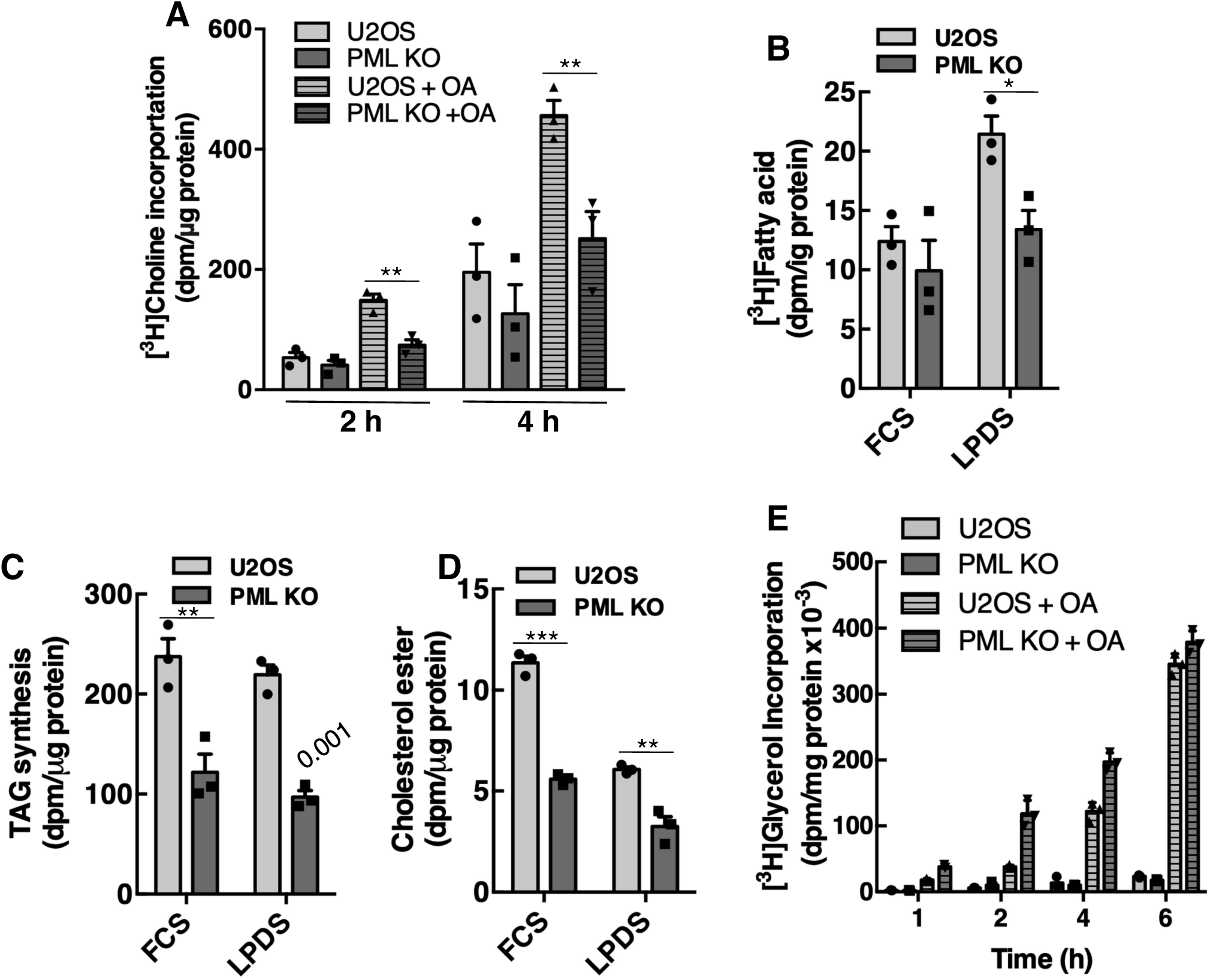
PML KO cells have reduced PC and TAG synthesis. A, Cells were cultured for 24 h in medium containing no addition or 300 μM oleate (OA), incubated in choline-free medium with [^3^H]choline (1 μCi/ml) for 2 or 4 h, and isotope incorporation into PC was quantified. Results are the mean and SD of 5 experiments. B, U2OS and PML KO cells were cultured in FCS or lipoprotein-deficient serum (LPDS) for 24 h, incubated with [^3^H]acetate for 4 h and incorporation into fatty acids was determined. Results are the mean and SD of 3 experiments. C and D, U2OS and PML KO cells were cultured in FCS or LPDS for 18 h prior to incubation with 100 μM [^3^H]oleate for 4 h to measure TAG and cholesterol ester (CE) synthesis. Results are the mean and SD of 3 experiments. E, U2OS and PML-KO cells were labelled with [^3^H]glycerol (2 μCi/ml) in the presence or absence of oleate (OA, 100 μM) for up to 6 h. [^3^H]Glycerol incorporation into TAG is the mean and SD of triplicate determinations from a representative experiment. Statistical comparisons by two-tailed t-test (\**p*<0.05; \*\**p*<0.01; \*\*\**p*<0.0001).

## DISCUSSION

The INM is contiguous with the outer nuclear membrane and cytoplasmic ER (Ungricht and Kutay, 2017) and thus could receive lipids by lateral diffusion from the ER (van Meer et al., 2008). However, the INM in yeast (Romanauska and Kohler, 2018) and mammals (Aitchison et al., 2015; Csaki et al., 2013; Haider et al., 2018; Peterson et al., 2011) harbours lipid biosynthetic enzymes and regulatory proteins suggestive of compartmentalized nuclear lipid synthesis. Analogous to the ER, the INM is the site for *de novo* biogenesis of nLDs in yeast (Romanauska and Kohler, 2018) whereas in hepatoma cells, nLDs are derived from luminal ER LDs that enter the nucleus after dissolution of the INM (Soltysik et al., 2019). PML expression, in particular the PML-II isoform, aids in nLD biogenesis at the INM by disrupting lamin-free regions of the INM to allow luminal LD release (Ohsaki et al., 2016). Ohsaki and colleagues also demonstrated that more than 70% of PML-containing structures in the nuclei of hepatoma cells were associated with the surface nLDs but their function is unknown. Here we demonstrate that these structures, which we refer to as LAP domains, are enriched in DAG and lipid biosynthetic enzymes but depleted of proteins that associate with canonical PML NBs. Further, LAP domains are not essential for nLD formation in U2OS cells but are necessary for the functional maturation of the LD into structures that recruit CCTα and Lipin1 for the regulation of PC and TAG synthesis.

Coincident with the appearance of LAP domains on nLDs in oleate-treated U2OS cells was the loss of canonical PML NB containing SUMO1, DAXX and SP100. While the expression of PML protein was unaffected by oleate and it appeared to transfer quantitatively to nLDs, the resultant LAP domains were poorly SUMOylated and depleted of PML NB resident proteins. Electron microscopy showed the surface of nLDs had the typical morphology of PML NB (Ohsaki et al., 2016), indicating that LAP domains are structurally related but have lost features that are required for PML NB assembly, notably SUMO1 and SP100. Importantly, since LAP domains are deficient in these normally constitutive protein components of this nuclear subdomain, by convention and definition they cannot be called PML NBs. By making this distinction, LAP domains are place within the continuum of PML-containing cellular structures that are already described in the literature with PML NBs at one extreme, followed by LAP domains, and at the far extreme, Mitotic Accumulation of PML Protein (MAPPs) that are completely devoid of DAXX, SP100 and SUMO1 (Dellaire et al., 2006).

PML KO U2OS cells had a 50-60% reduction in the number of nLDs, a shift to smaller size and reduced association of CCTα, Lipin1 and DAG. However, large CCTα-positive nLDs in PML KO cells were only reduced by 50% indicating other mechanisms for nLD biogenesis. Microsomal triglyceride transfer protein inhibitors did not affect nLD formation in U2OS cells suggesting luminal LDs may not be precursors (Soltysik et al., 2019). However, PML-negative nLDs were observed in close proximity to the NE suggesting nLD could originate from LD precursors and/or *de novo* lipid synthesis at the INM. A clue to the potential mechanisms for nLD biogenesis comes from high resolution imaging of the polarized localization of LAP domains on nLDs. PML-II patches on the INM have been proposed as sites where nLD emerge from the INM (Ohsaki et al., 2016). 3D imaging revealed that these LAP domain patches persist on small and large nLDs and likely represent the surface at which the nascent nLD acquired PML proteins as it emerged from the INM. The subsequent maturation of the nLD could be facilitated by the recruitment and activation of CCTα and Lipin1 on the exposed surfaces of the LD monolayer and accompanying increase in PC and TAG synthesis. The emergence of nLDs from the INM could also involve the CCTα M-domain, which is capable of inducing positive membrane curvature and expansion of the NR (Lagace and Ridgway, 2005; Taneva et al., 2012). In this scenario, translocation of CCTα to the INM could induce NR formation and destabilization of the bilayer to facilitate the release of nascent nLDs. The presence of large CCTα-positive nLDs in PML KO cells supports this concept.

Since PML KO prevented the increase in PC synthesis and reduced CCTα-positive nLDs, we conclude that LAP domains are essential to activate PC synthesis in response to prolonged exposure to fatty acids. Analysis of CCTα mutants showed that association with nLDs was mediated by basic and acidic residues in the M-domain and negatively regulated by phosphorylation of the P-domain. These domains also regulate CCTα association with membrane *in vitro*, demonstrating that interactions with nLDs are lipid-mediated. Thus LAP domains promote the formation of nLDs on to which CCTα can translocate and activate PC synthesis. To identify lipids that promote CCTα interaction with nLDs we initially focused on DAG, which induces packing defects in membranes into which the CCTα M-domain can insert (Cornell, 2016; Cornell and Ridgway, 2015). PML expression was positively correlated with large nLDs that were enriched in DAG. However, DAG and CCTα were not preferentially co-localized in nLDs nor was their relative distribution in nLDs altered by loss of PML expression. While the exclusion of CCTα and the biosensor on nLDs could be due to competition for DAG, it is more likely that factors such as surface curvature, enrichment in anionic lipids or a high PE/PC ratio promote the association of CCTα. For instance, the high PE/PC ratio in insect cLDs relative to human cLDs (Jones et al., 1992; Tauchi-Sato et al., 2002) could be responsible for nuclear export and localization of CCT1 and CCTα on cLDs during oleate loading in *Drosophila* S2 cells (Krahmer et al., 2011).

The striking PML-dependent redistribution of DAG between nLDs and cLDs was linked to the nuclear PA phosphatase Lipin1α, which co-localized with DAG on nLDs in oleate-treated U2OS and Caco2 cells. Since small nLDs in PML KO cells were relatively devoid of both Lipin1α and DAG, Lipin1α appears to provide DAG for nuclear TAG synthesis by diacylglycerol acyltransferase 2, which was localized to nLDs when ectopically expressed in Huh7 cells (Ohsaki et al., 2016). The factors mediating the selective association of Lipin1 with nLDs are uncertain. PA, which promotes Lipin1 association with membranes (Ren et al., 2010), was not detected on nLDs or cLDs with a biosensor suggesting it is below the detection threshold or the biosensor is not sufficiently PA-specific, as was suggested previously (Horchani et al., 2014). Lipin1α was not detected on cLDs and associated with accumulation of DAG on cLDs in PML KO cells, which could reflect a partial block in TAG synthesis or increased TAG hydrolysis.

Metabolic labelling experiments support the concept that LAP domains and nLDs are sites for TAG synthesis and regulation. PML KO cells had significantly reduced [^3^H]oleate incorporation into TAG, reduced nLDs, and increased levels of DAG in cLDs, indicative of a block in nuclear and cytoplasmic TAG synthesis. However, PML KO did not inhibit *de novo* synthesis TAG from [^3^H]glycerol suggesting a more complex role in oleate uptake and utilization. Prior studies have indicated complex, tissue-specific roles for PML in fatty acid metabolism. PML NB-dependent activation of peroxisome proliferator-activated receptor (PPAR) γ co-activator-1α (PGC-1α), PPAR signaling and fatty acid β-oxidation provided a growth advantage to breast cancer cells (Carracedo et al., 2012) and controlled asymmetric division and maintenance of hematopoietic stem cells (Ito et al., 2012). In contrast, PML KO mice had tissue-specific enhancement of both fatty acid β-oxidation and synthesis, increased metabolic rate and resistance to diet-induced obesity (Cheng et al., 2013). Together with these findings, our results suggest that LAP domains associated with nLDs could have a multi-facetted role in lipid homeostasis involving the recruitment of CCTα and Lipin1 to promote PC and TAG synthesis and storage as well as Lipin1 regulation of a PGC-1α/PPAR signaling pathways that regulate fatty acid uptake, storage and oxidation (Finck et al., 2006).

## ACKNOWLEDGEMENTS

We would like to thank Romain Laine and Ricardo Henriques (University College London/Francis Crick Institute, UK) for providing guidance on SRRF imaging and optimization. This work was funded by a Project Grant to GD and NDR from the Canadian Institutes of Health Research (PJT62390) and the Bernard and Winnifred Lockwood Endowment for Research. JL was supported by a scholarship from the Nova Scotia Health Research Foundation. GD and NDR are Senior Scientists of the Beatrice Hunter Cancer Research Institute (BHCRI).

## MATERIALS AND METHODS

### Cell culture

U2OS (ATCC HTB-96) and CaCo2 (ATCC HTB-37) cells were cultured in DMEM containing 10% FBS (medium A) at 37°C in a 5% CO2 atmosphere. CRISPR/Cas9 methodology was used to knockout the expression of all PML isoforms in U2OS cells (Attwood et al., 2019). Unlabeled and [^3^H]oleate/BSA complexes (6:1, mol/mol, 12 mM oleate stock solutions) were prepared as described (Goldstein, 1983).

### Plasmid transfection

Cells were transfected with plasmids encoding murine Lipin1α-V5 and Lipin1β-V5 (Bou Khalil et al., 2009) (Zemin Yao, University of Ottawa), EGFP-PML isoforms (Bischof et al., 2002) (Oliver Bischof, Institute Pasteur), GFP-C1(2)δ (Tobias Meyer, Addgene plasmid #21216), GFP-nes-2xPABP (Sergio Grinstein, University of Toronto), CCTα-16SA and -16SE (P-domain phosphorylation mutants) (Wang and Kent, 1995), CCTα-3EQ and -8KQ (M-domain mutants (Gehrig et al., 2008; Johnson et al., 2003) using Lipofectamine 2000 (2 μl/μg DNA) according to manufacturer’s instructions (Life Technologies, Burlington, ON). pGFP-C1(2)δ-2NLS was prepared by subcloning a x2 nuclear localization signal (NLS) cassette into the BsrGI-AflII site of pGFP-C1(2)δ. Experiments were initiated 24-36 h after plasmid transfection.

### Immunoblotting

Cells were lysed in SDS buffer (12.5% SDS, 30 mm Tris-HCl, 12.5% glycerol, and 0.01% bromophenol blue, pH 6.8), heated at 95°C for 5 min. Lysates were separated by SDS-PAGE, transferred to nitrocellulose membranes and incubated in TBS (20 mM Tris-HCl [pH 7.4] and 500 mM NaCl):Odyssey blocking (5:1, v/v) for 1h. Nitrocellulose was blotted using primary antibodies against human CCTα (Morton et al., 2013), PML (rabbit polyclonal A301–167A, Bethyl Laboratories), V5 monoclonal (MCA-1360, BioRad), or β-actin (mouse monoclonal AC15, Sigma-Aldrich). Proteins were visualized with goat anti-mouse or goat anti-rabbit IRDye-800 or - 680 secondary antibodies (LI-COR Biosciences) using an Odyssey Imaging System and application software (v3.0).

### Analysis of PC synthesis using [^3^H]choline incorporation

U2OS and PML KO cells were incubated in the presence or absence of 400 μM oleate/BSA for 24 h. Cells were rinsed twice with choline-free medium A and then incubated with choline-free medium A containing [^3^H]choline (2 μCi/ml) for 2 and 4 h. After isotope labeling, cells were rinsed with cold PBS, harvested in methanol:water (5:4, v/v), [^3^H]PC was extracted in chloroform/methanol, and the radioactivity quantified by liquid scintillation counting and normalized to total cellular protein (Storey et al., 1997).

### Analysis of fatty acid and TAG synthesis

U2OS and PML KO cells were incubated with 2.5 μCi/ml [^3^H]acetate for 4 h to measure fatty acid synthesis (Brown et al., 1978). Briefly, lipid extracts from cells were saponified in ethanol and 50% potassium hydroxide (w/v) for 1 h at 60°C and extracted with hexane. The hydrolysate was then acidified with HCl (pH <3) and radiolabeled fatty acids were extracted with hexane and quantified by liquid scintillation counting. Incorporation of [^3^H]acetate into fatty acids was normalized to total cellular protein.

TAG and cholesterol ester (CE) synthesis from exogenous fatty acid was determined by incubating cells with 100 μM [^3^H]oleate/BSA. After 4h, cells were rinsed twice with cold 150 mM NaCl, 50 mM Tris-HCl (pH 7.4) with 2 mg/ml BSA and once with the same buffer without BSA. [^3^H]Oleate-labeled lipids were extracted from dishes with hexane:isopropanol (3:2, v/v). Samples were dried under nitrogen, resolved by thin-layer chromatography in hexane:diethyl ether:acetic acid (90:30:1, v/v) and radioactivity in [^3^H]TAG and [^3^H]CE was quantified by scintillation counting and normalized to total cellular protein. *De novo* TAG synthesis was determined by incubating cells with or without 400 μM oleate/BSA in the presence of 2 μCi/ml [^3^H]glycerol. Cells were rinsed with cold PBS and radioactive lipids were extracted and quantified as described above.

### Immunofluorescence microscopy

Cells cultured on glass coverslips were fixed with 4% (w/v) paraformaldehyde and permeabilized for 10 min with 0.1% (w/v) Triton X-100 at room temperature. Coverslips were incubated with primary antibodies against CCTα, PML (monoclonal E-11, Santa Cruz), PML (rabbit polyclonal A301–167A, Bethyl Laboratories), LMNA/C (monoclonal 4C11, Cell Signaling), emerin (polyclonal FL-254, Santa Cruz), DAXX (polyclonal D7810, Sigma), SP100 (polyclonal PA5-53602, Invitrogen), SUMO1 (rabbit monoclonal Y299, Ab32058, Abcam) or V5 in PBS containing 1% (w/v) BSA. This was followed by secondary AlexFluor488-, 594- and 647- conjugated goat anti-rabbit or anti-mouse secondary antibodies (Thermo Fisher Scientific). To visualize nuclear and cytoplasmic LDs, BODIPY 493/503 or LipidTox Red was diluted 1:1,000 or 1:500, respectively, and incubated with the secondary antibodies. Coverslips were mounted on glass slides in Mowiol 4-88, and confocal imaging was performed using a Zeiss LSM510 or LSM710 laser-scanning confocal microscope with a Plan-Apochromat 100× (1.4 NA) oil immersion objective. LD cross sectional area in confocal images was quantified using ImageJ software (v1.47, National Institutes of Health). Images were converted to 8-bit, the threshold was adjusted, and the “analyze particle” command was used to exclude on edges. The percent distribution of LDs within a binned area group was quantified.

For super resolution radial fluctuation (SRRF) imaging (Gustafsson et al., 2016), immunostained cells (described above) were observed using a 100 X Plan-Apochromat (1.46 NA) oil immersion objective lens (Zeiss) by wide-field imaging on a Marianis microscope (Intelligent Imaging Innovations, 3i) based on a Zeiss Axio Cell Observer equipped with LED-based illumination via a SPECTRA III Light Engine (Lumencor), and a Prime BSI back-illuminated scientific complementary metal–oxide–semiconductor (sCMOS) camera (Teledyne Photometrics). This imaging configuration resulted in an effective pixel size of 65 x 65 nm in the captured images. To generate super resolution images, 100 wide-field images were captured at 10 ms/frame using Slidebook 6 software and then exported to ImageJ (version 1.52, NIH) in a 16-bit Open Microscopy Environment-Tagged Image File Format (OME-TIFF). Image processing used a custom SRRF algorithm (NanoJ-LiveSRRF, available on request from Ricardo Henriques, University College London/Francis Crick Institute, UK) with the following settings: radius 3; sensitivity 2; magnification 4; average temporal analysis; and with intensity weighting and both vibration and macro-pixel correction turned on. NanoJ-LiveSRRF is the newest implementation of NanoJ-SRRF within the ImageJ software, available upon request as above. However, NanoJ-SRRF is already freely available (Gustafsson et al., 2016).

For confocal imaging and 3D volume rendering, cells were immunostained as above and imaged using a 100 X Plan-Apochromat (1.46 NA) oil immersion objective lens (Zeiss) by spinning-disk confocal microscopy on a Marianis microscope (Intelligent Imaging Innovations, 3i) based on a Zeiss Axio Cell Observer equipped with Yokagawa CSU-X1 spinning-disk unit, four laser lines (405, 488, 560 and 640 nm) and a Prime95B (Teledyne/Photometrics). 3D volume rendering of confocal image stacks was done using Slidebook 6 software (3i).

