## Supplemental figures 1-9 for "Lipid-associated PML domains regulate CCTα, Lipin1 and lipid homeostasis"

### Lipid-Associated PML (LAP) domains on nuclear lipid droplets harbour CCTα and Lipin1 and regulate lipid homeostasis

### Supplemental Figure S1

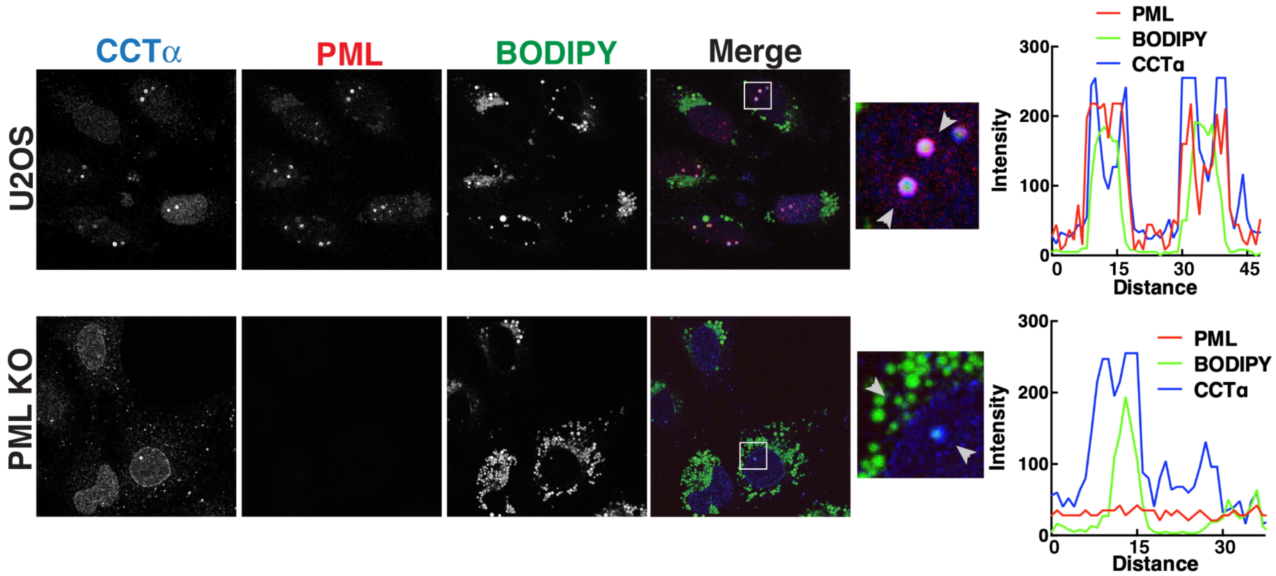


**Supplemental Figure S1**. **Co-localization of PML and CCTα on nLDs.** U2OS and PML KO cells were cultured in oleate (400 μM) for 24 h prior to fixation and immunostaining for CCTα (blue) and PML (red) and imaging by confocal microscopy. LDs were visualized with BODIPY 493/503 (green). Arrows indicate the region selection for RGB line plots showing the co-localization of PML and CCTα on the surface of nLDs and the NE.

Supplemental Figure S2

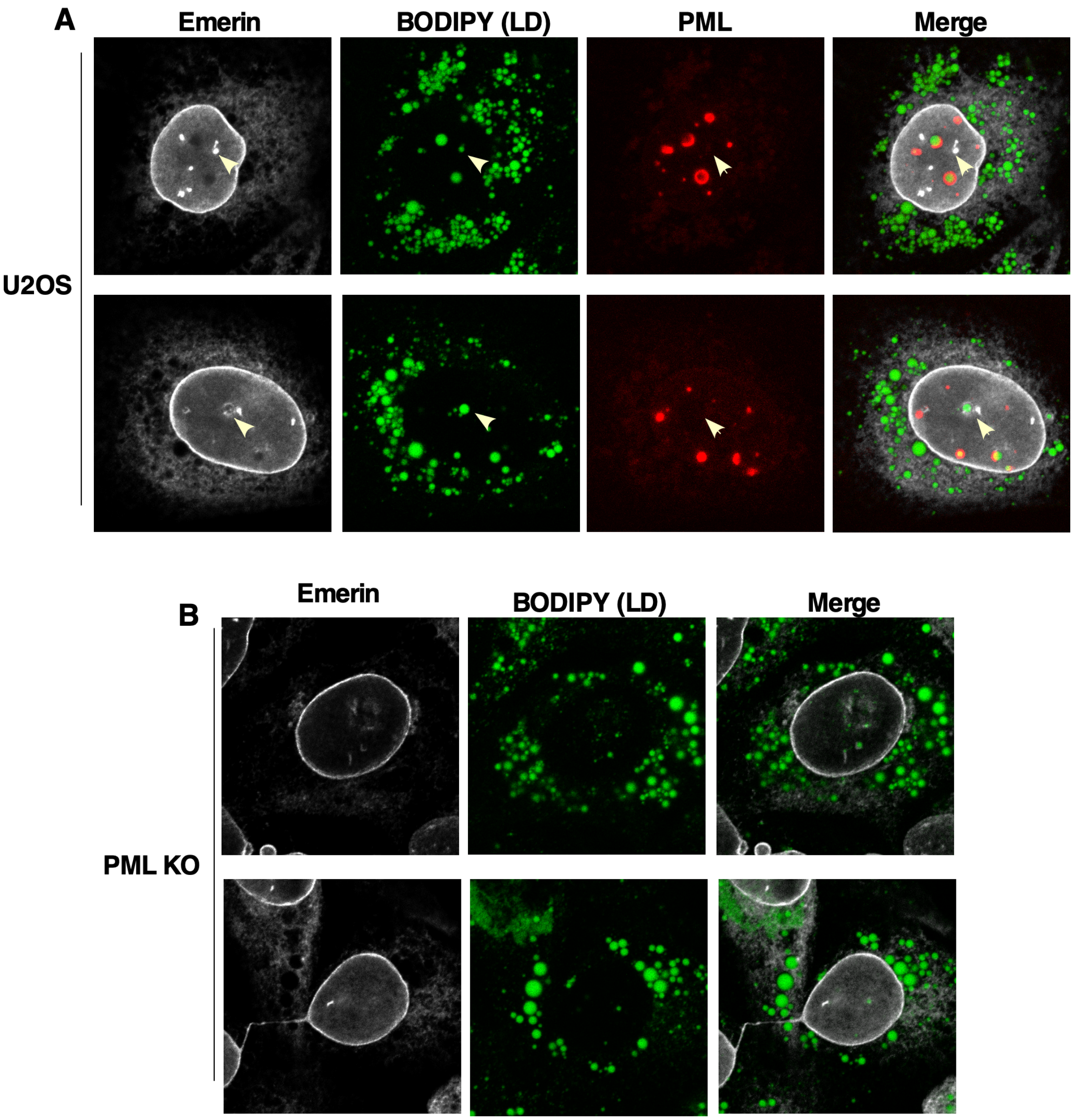


**Supplemental Figure S2**. **PML-negative nLDs associate with the nuclear envelope and nucleoplasmic reticulum.** A, U2OS cells were cultured in oleate (400 µM) for 24 h prior to fixation and immunostaining for emerin (white) and PML (red) and imaging by confocal microscopy. LDs were visualized with BODIPY 493/503 (green). Arrows indicate the position of PML-negative nLDs that are associated with emerin-positive nucleoplasmic reticulum. B, PML KO cells were treated as described above and immunostained for emerin (white) and LDs were visualized with BODIPY (green).

Supplemental Figure S3

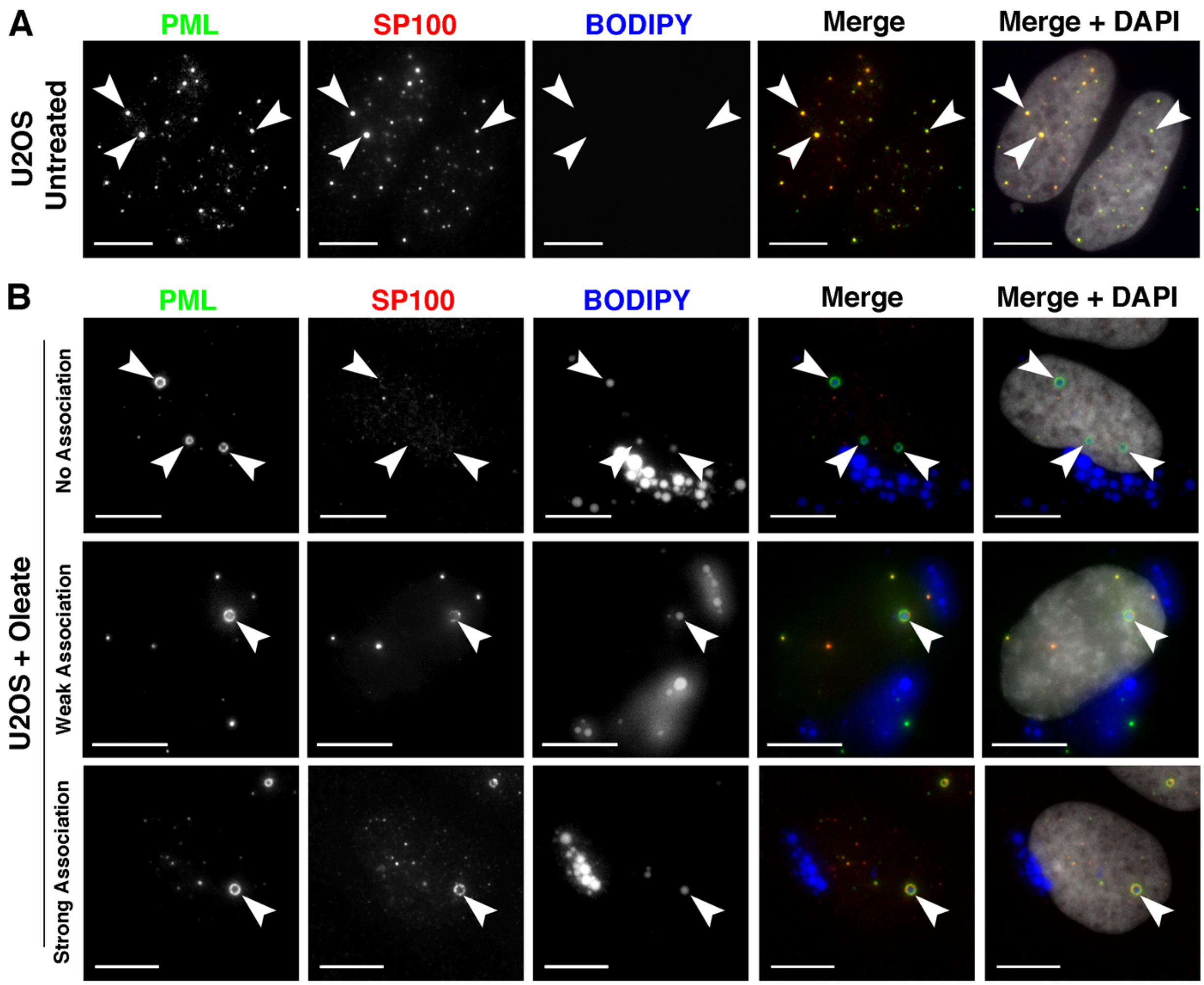


**Supplemental Figure S3**. **LAP domains are deficient in SP100**. U2OS cells were untreated (panel A) or treated with oleate (400 µM) for 24 h (panel B) prior to fixation and immunostaining for SP100 (red) and PML (green) and imaging by confocal microscopy. LDs were visualized with BODIPY 493/503 (blue) (bar, 10µm). Images were captured as described in the legend to Fig. 1.

Supplemental Figure S4

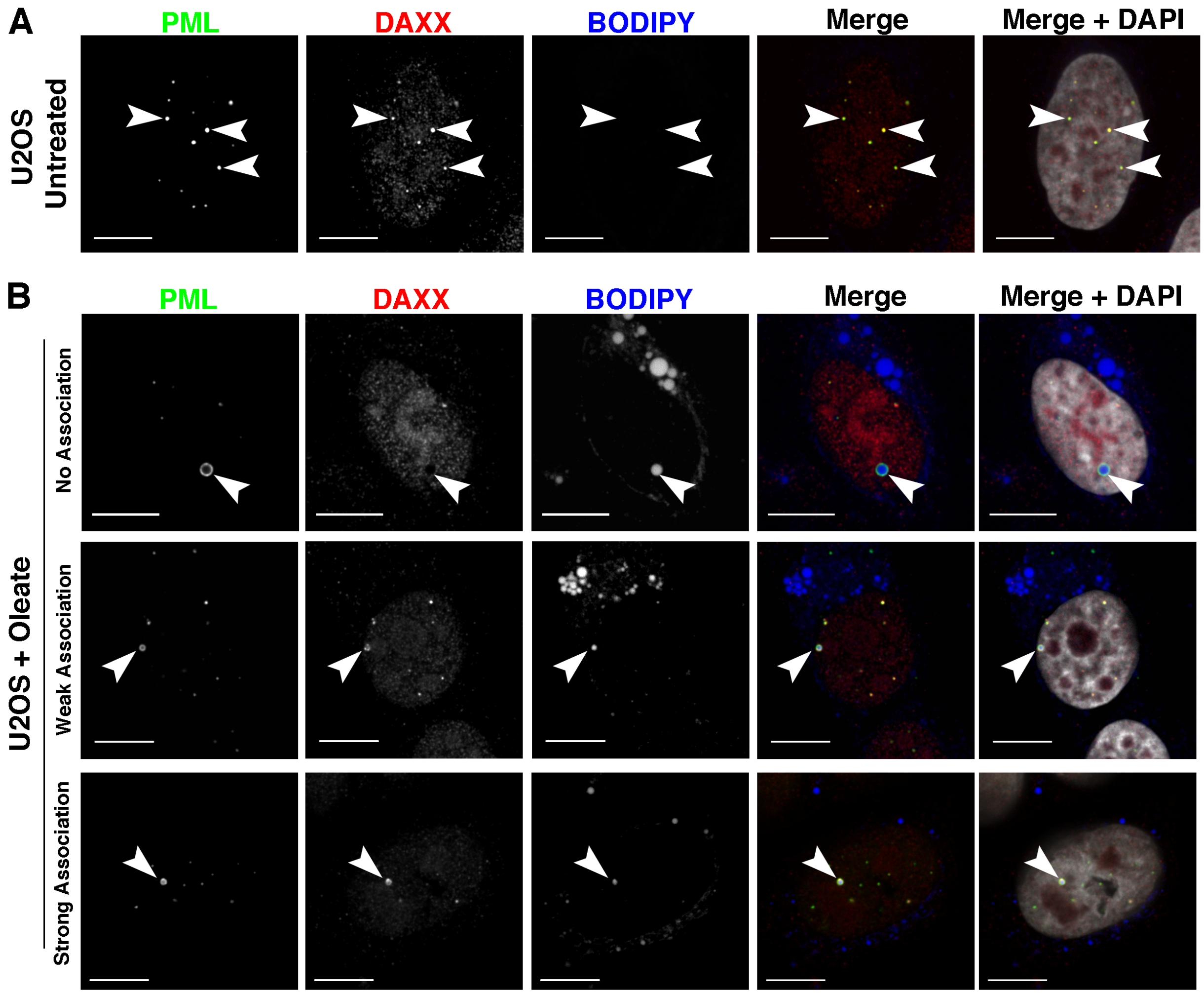


**Supplemental Figure S4**. **LAP domains are deficient in DAXX.** U2OS cells were untreated (panel A) or treated with oleate (300 μM) for 24 h (panel B) prior to fixation and immunostaining for DAXX (red) and PML (green) and imaging by confocal microscopy. LDs were visualized with BODIPY (blue) (bar, 10μm).

Supplemental Figure S5

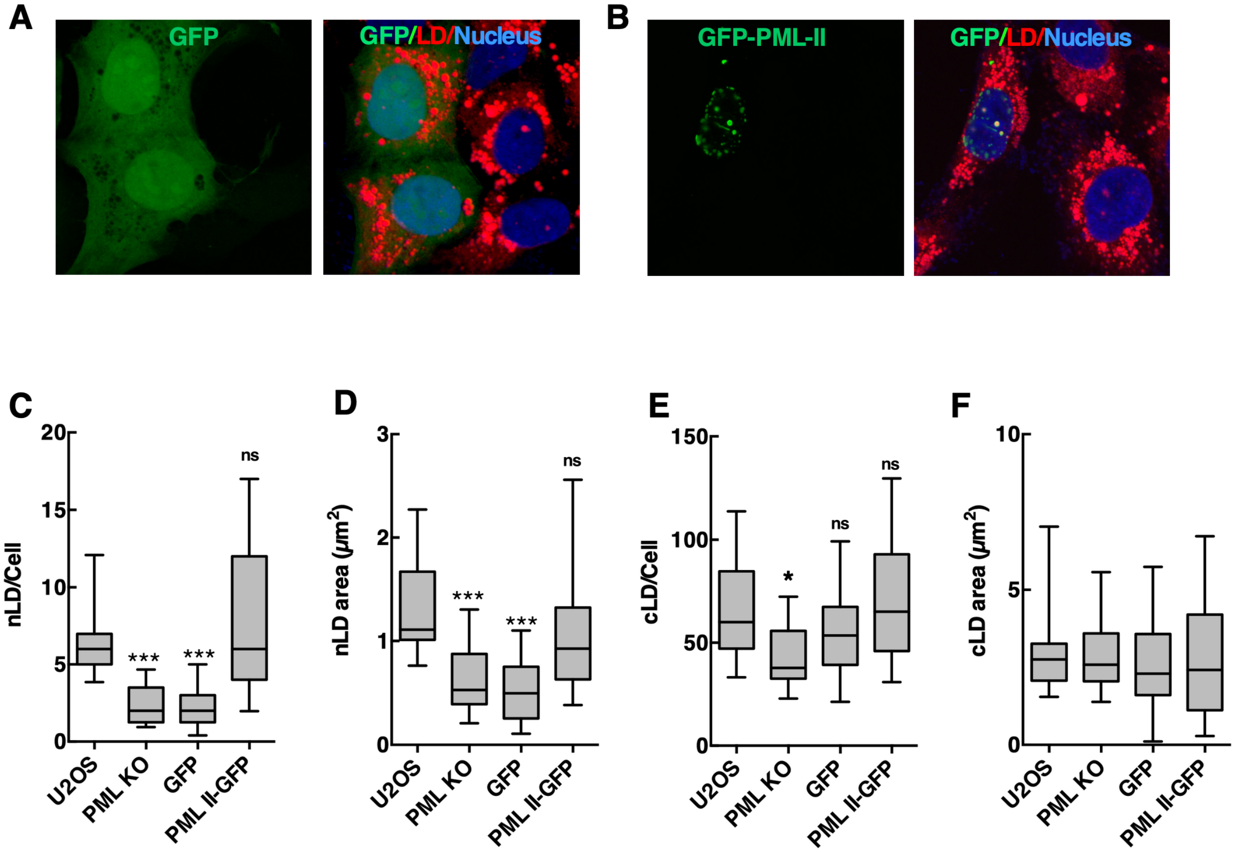


**Supplemental Figure. S5**. **Expression of GFP-PML-II restores nLD in PML KO cells**. PML-KO cells were transiently transfected with pGFP-N1 (panel A) or pGFP-PML-II (panel B) for 24 h followed by incubation with 400 µM oleate for 24 h. Nuclei were stained with Hoechst 33342 (blue) and LDs were visualized with LipidTox Red and imaged by confocal microscopy. C and D, quantification of nLD number and cross-sectional area in cells. E and F, quantification of cLD number and cross-sectional area in cells. Results in panels C-F are presented as box and whisker plots showing the mean and $5^{th}$-to-$95^{th}$ percentile for analysis of 50-100 cells from 3 separate experiments involving 25-50 cells. Significance was determined by one-way ANOVA and Tukey's multiple comparison (ns, not significant; (*$p<0.05$; ***$p<0.0001$).

Supplemental Figure S6

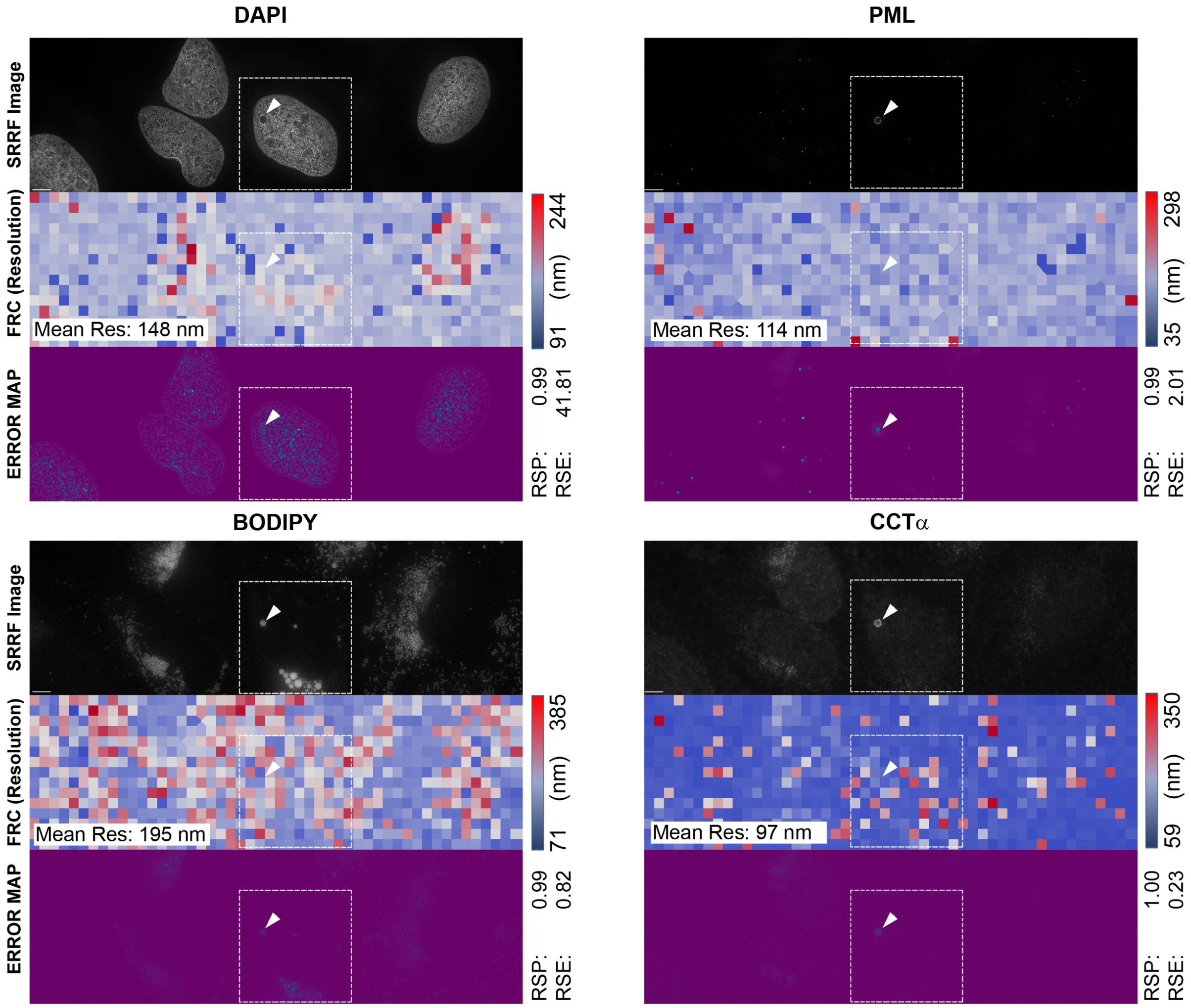


**Supplemental Figure S6. Uncropped Super-Resolution Radial Fluctuation (SRRF) Images from Figure 4 and Error and Resolution Analysis using NanoJ-SQUIRREL (Culley et al., 2018).** The original SRRF image before cropping from Figure 4 is shown for DAPI, PML (AlexaFluor-647), BODIPY 493/503 and CCTα (AlexaFluor-568) fluorescent micrographs in the top row of each image grouping. The middle row in each grouping represents the resolution estimate divided in ~750 blocks by Fourier Ring Correlation (FRC) analysis (mean estimated resolution is shown in the inset at the left). The bottom row represents an Error Map based on the Resolution Scale Pearson (RSP) Correlation and the Resolution Scale Error (RSE).

Supplemental Figure S7

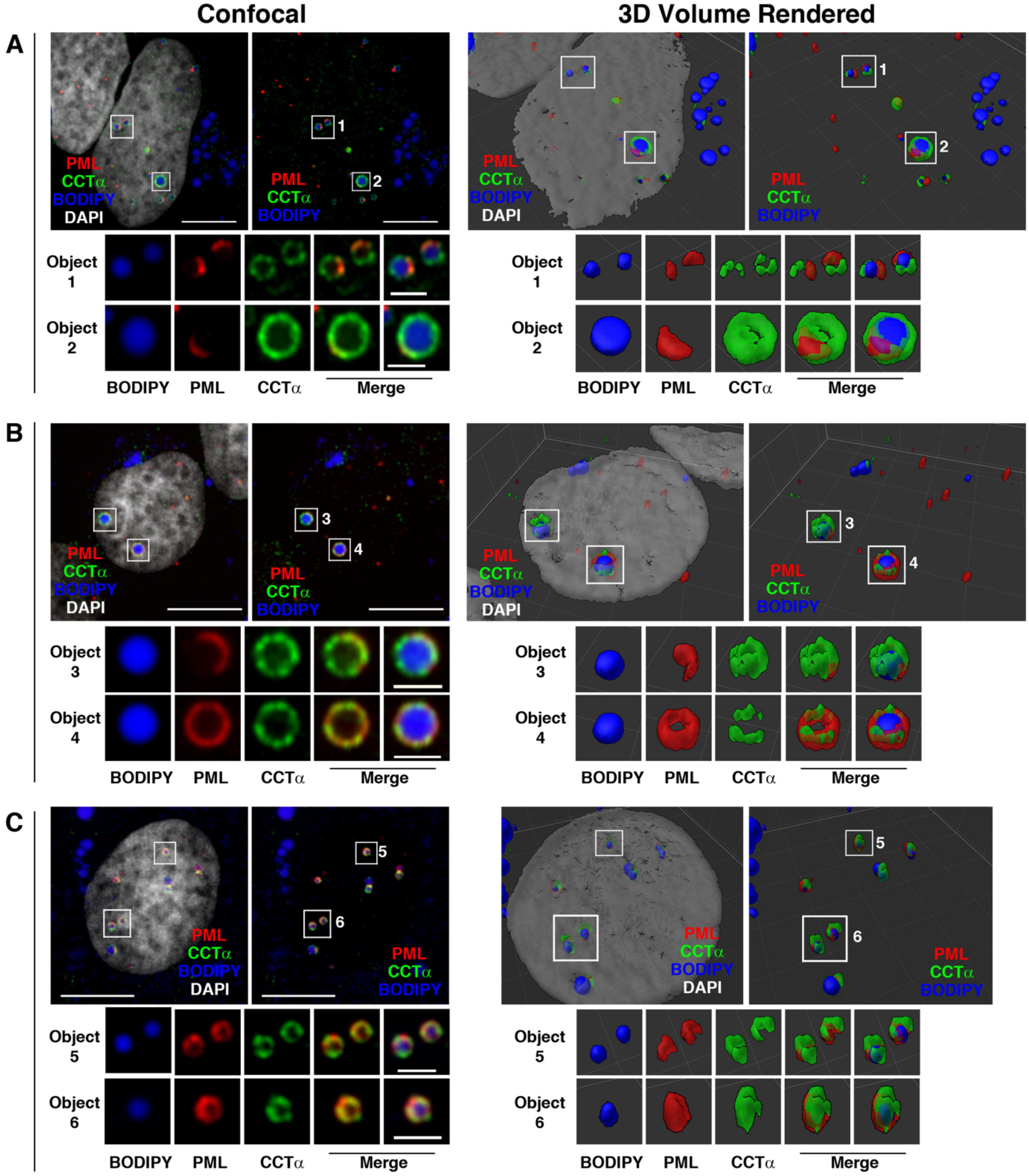


**Supplemental Figure S7. Spinning disk confocal 3D imaging of CCTα and PML on nLDs**. U2OS cells were treated with oleate for 24 h prior to fixation and immunostaining for CCTα (green) and PML (red). nLDs were localized based on BODIPY 493/503 as described in the legend to Fig. 4. Spinning disk confocal sections and full 3D rendering (Slidebook) are shown for 3 representative nuclei (panels A, B and C) and 8 nLDs are enlarged (objects 1-6) to better show the relationship between PML and CCTα on nLDs. DNA was visualized with DAPI. The scale bar on confocal images is 10 μm (inset is 2 μm) and the grid on the 3D render is 5 μm.

Supplemental Figure S8

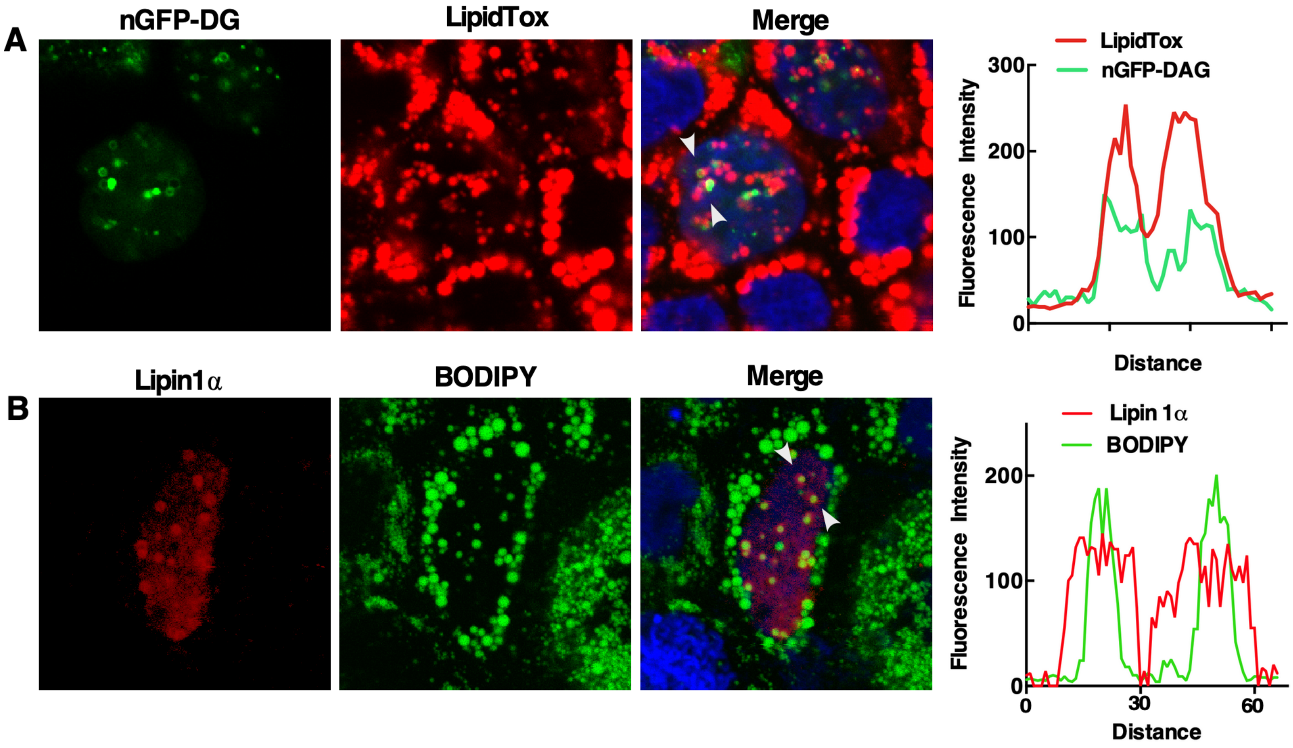


**Supplemental Figure S8**. **Detection of DAG and Lipin1α in the nLD of oleate-treated Caco2 cells.** A, Caco2 cells transiently expressing nGFP-DAG were treated with oleate/BSA (400 μM) for 24 h prior to fixation for confocal microscopy. LDs were visualized with LipidTox Red. B, Caco2 cells expressing Lipin1α-V5 (red) were treated as described above. LDs were visualized with BODIPY493/503 (green). In panels A and B, nuclei were stained with Hoechst 33258 (blue). Arrows indicate the region selected for associated RGB line plots showing DAG-GFP or Lipin1α on the surface of nLD.

Supplemental Figure S9

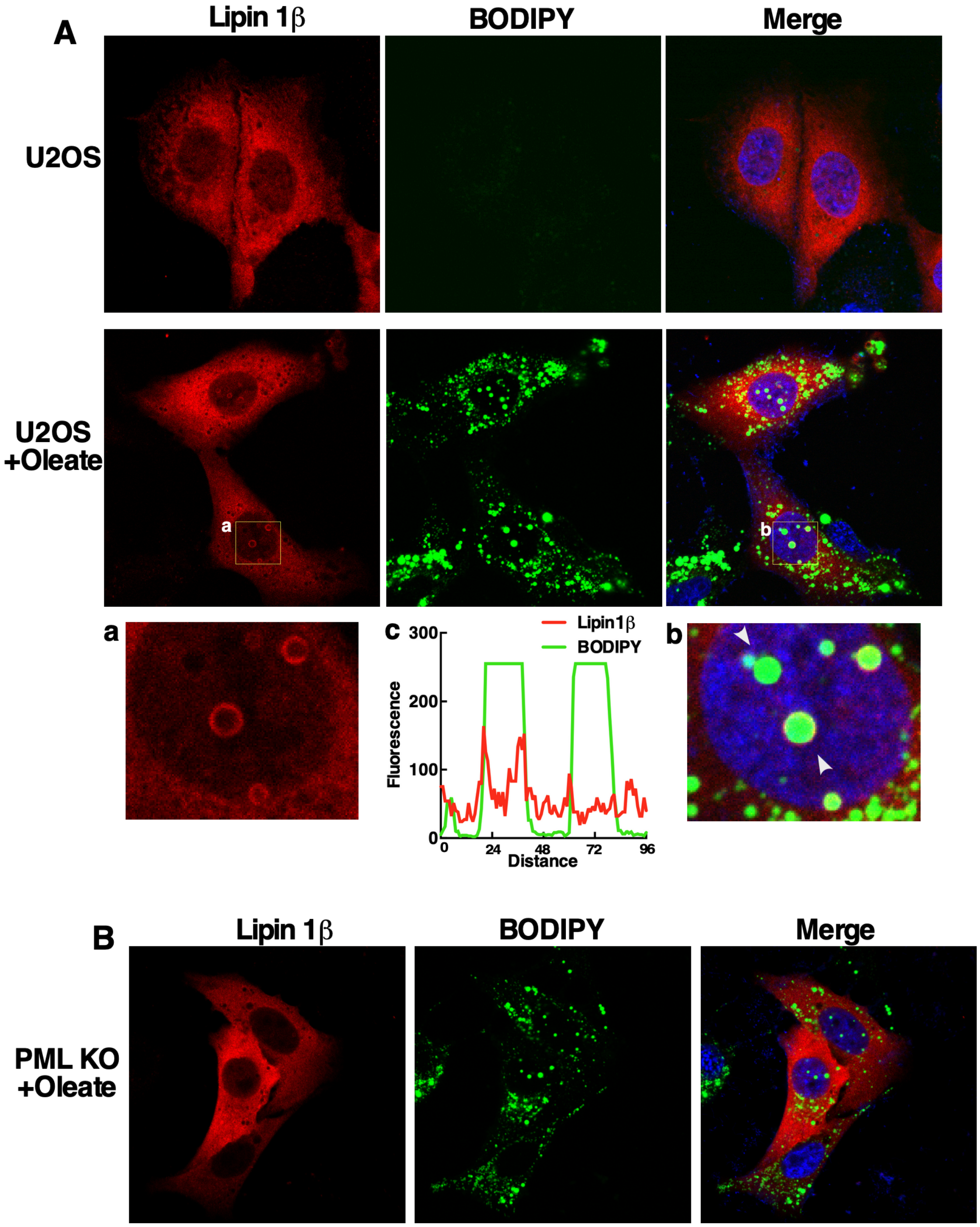


**Supplemental Figure S9**. **Lipin1β association with nLDs is dependent on PML expression**. A, U2OS cells transiently expressing Lipin1β-V5 were untreated or incubated with oleate (400 µM) for 24 h prior to fixation and immunostaining with a V5 monoclonal antibody (red) and imaged by confocal microscopy. Selected regions from Lipin1β (a) and merge (b) images of oleate-treated cells show the association of Lipin1β with nLD stained with BODIPY 493/503 (green). Arrows indicate the region selected for RGB line plots (c) showing Lipin1β is on the surface of nLD. B, Transiently expressed Lipin1β in oleate-treated PML KO cells (as described in panel A) was not detected on the surface of nLDs.

Supplemental Figure S10

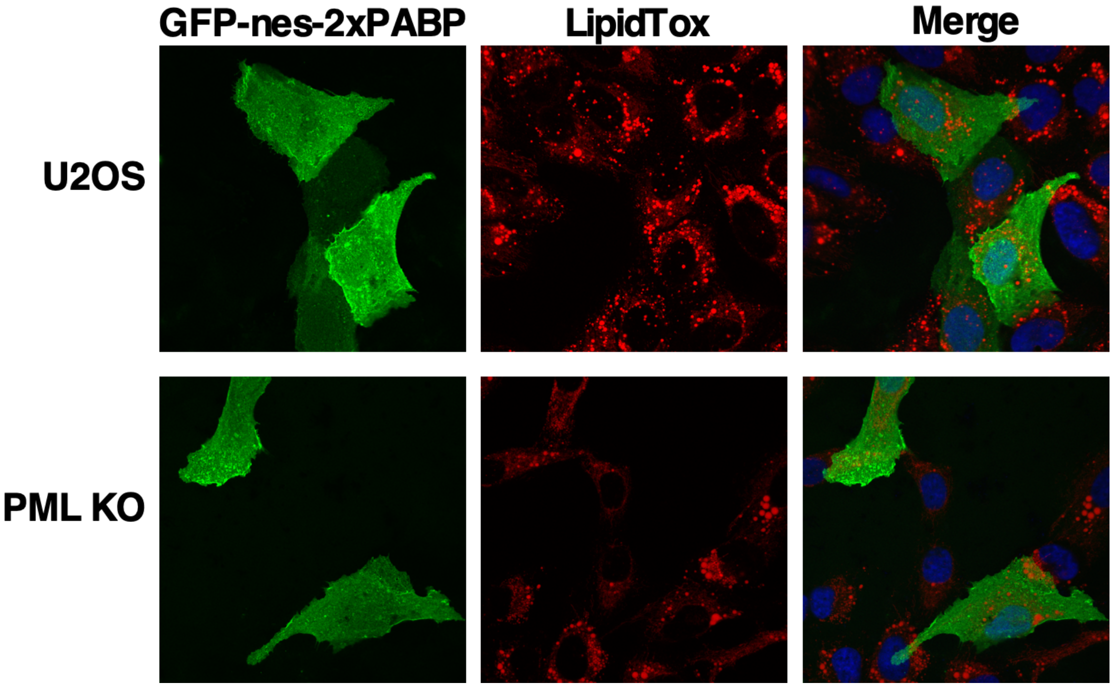


**Supplemental Figure. S10. Phosphatidic acid is not detected on nLD or cLD.** U2OS and PML KO cells transiently expressing the PA biosensor GFP-nes-2xPABP were cultured in media with oleate (300 μM) for 24 h prior to fixation and confocal microscopy. LDs were visualized with LipidTox Red.
